# Modeling Human Visuomotor Adaptation with a Disturbance Observer Framework

**DOI:** 10.1101/2025.09.01.673530

**Authors:** Gaurav Sharma, Bernard Marius ’t Hart, Jean-Jacques Orban de Xivry, Denise Y.P. Henriques, Mireille E. Broucke

## Abstract

A fundamental problem of visuomotor adaptation research is to understand how the brain is capable to asymptotically remove a predictable exogenous disturbance from a visual error signal using limited sensor information by re-calibration of hand movement. From a control theory perspective, the most striking aspect of this problem is that it falls squarely in the realm of the internal model principle of control theory. Despite this fact, the relationship between the internal model principle and models of visuomotor adaptation is currently not well developed.

This paper aims to close this gap by proposing an abstract discrete-time state space model of visuomotor adaptation based on the internal model principle. The proposed *DO Model*, a metonym for its most important component, a disturbance observer, addresses key modeling requirements: modular architecture, physically relevant signals, parameters tied to atomic behaviors, and capacity for abstraction. The two main computational modules are a disturbance observer, a recently developed class of internal models, and a feedforward system that learns from the disturbance observer to improve feedforward motor commands.

**Author summary:** A central research challenge in the field of motor learning is to disentangle the components of motor learning [1–8]. The main hypothesis explored in this paper is that visuomotor rotation experiments in which participants are given distinct instructions to either move the hand or the cursor to the target offer a platform to explore two components of motor learning arising from different computations and possibly different brain regions. The two computations correspond to a *disturbance observer* whose role is to detect and then eliminate environmental perturbations and a *feedforward system* whose role is to learn from the disturbance observer to improve feedforward motor commands.

## 1 Introduction

A central challenge in the area of modeling visuomotor adaptation is to achieve a behavioral discrete-time state space model that clarifies the precise computations driving this brain process. The model should be sufficiently abstract to avoid low-level neural computations, yet be sufficiently refined to capture the essence of the brain computations driving this process. Developing such a model will be a multi-step undertaking, involving successive generations of increasingly accurate behavioral models. This paper aims to take a step along this path. The following requirements should be addressed.

i. **Modular Control Architecture**. The model must identify all major (high-level) computational modules involved in forming the motor command in the brain for performing visuomotor adaptation. The functional role of each computational module must be explicitly motivated. The flow of information and the control architecture of the model must be explicitly represented. Relevant timescales involved in generating the motor command must be identifiable. Silent computations as well as concurrent computations should be clarified. The model must be closed-loop stable (a stability analysis is not performed in this paper).
ii. **Measurements and States**. Measurements arriving in the brain from the body’s sensors must be biologically plausible based on what is know about visuomotor adaptation and related functions such as saccade adaptation. Brain states (states originating in the brain) and physical states (such as the hand angle) must be clearly distinguished from each other. The role of each brain state must be explicitly justified. A state cannot be an abstract signal whose origin is unknown.
iii. **Parameters**. Parameters should have a clear physical meaning, each parameter should be associated with an atomic behavior to the extent possible, and there should be explicit procedures to identify each parameter. Curve fitting multiple parameters to data should be avoided.
iv. **Abstraction**. Computational modules corresponding to processes evolving in the world must either adhere to the laws of physics or be plausible abstractions thereof. The model should avoid low-level models of individual neurons or neural networks. It should avoid details of the physics of the arm. On the other hand, there should be a clear placeholder where the modeler may insert such additional details.
v. **Expressivity and Predictive Ability**. The model should explain a breath of experimental phenomena in visuomotor adaptation and also should be capable to make predictions about new behaviors.

We propose a discrete-time behavioral state-space model of visuomotor adaptation designed to address this modeling challenge. The *DO Model*, whose name is a metonym for its most important component, a disturbance observer, accounts for both the experimental findings reported here and phenomena observed in prior studies. The model architecture (Fig 1) consists of three control components: (i) a fast-reacting error feedback process, which aligns with single-trial learning effects; (ii) a disturbance observer, which estimates persistent perturbations in the visual error signal and drives their asymptotic elimination; and (iii) a slower feedforward learning system that gradually updates internal control commands through a process of learning transfer from the disturbance observer.

**Fig 1.**
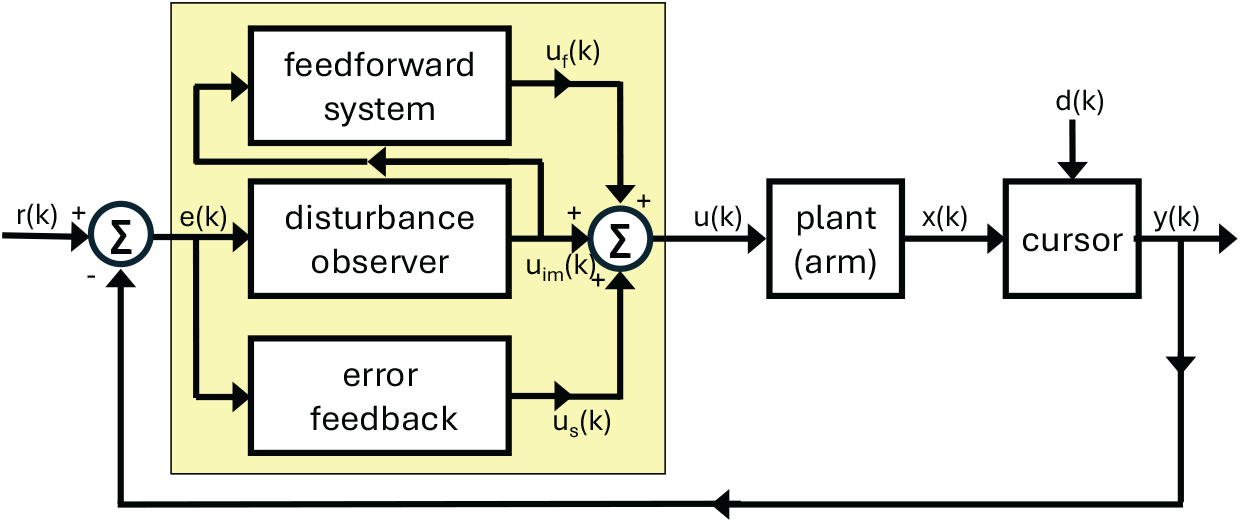
Block diagram of the DO model during Learn trials.

The disturbance observer and feedforward system are dynamically coupled; however, the dominant mode of interaction is a cascade architecture in which the disturbance observer continuously influences the feedforward learner. Additionally, the model includes the possibility of a phase switch, representing a transition between the actively expressed computational module or strategy, consistent with observations of context-dependent changes in behavior.

The disturbance observer functions as a specialized internal model that isolates and compensates for persistent exogenous disturbances affecting the sensory error signal. Disturbance observers have not, to our knowledge, been systematically applied within sensorimotor modeling. A central aim of this paper is to demonstrate their utility for explaining key features of visuomotor adaptation.

The second major component, the feedforward system, underlies the ability to execute accurate reaching movements in the absence of visual feedback or under instructions to “move the hand to the target.” It updates internal motor commands based on sustained discrepancies identified by the disturbance observer — effectively learning from evidence of persistent miscalibration. This process unfolds over a medium to long timescale and supports adaptation beyond immediate error-based corrections. Thus, the feedforward system is conceived as a motor system that gradually adapts to the presence of persistent activity in the disturbance observer, through a process of of *learning transfer, consolidation of motor memory*, or *transfer of memory trace* [9–14]. This part of the model has been inspired by modeling studies of the floccular complex and its relationship to the medial vestibular nucleus [15, 16].

The inclusion of both the disturbance observer (driving computations when a cursor is visible and the participant is instructed to move the cursor to the target) and a feedforward system (driving computations when there is no cursor or the instructions are to move the hand to the target) means that the DO model can also account for so-called *phase switches*, switches in computational streams in the brain, a capability concurring with experimental studies on rapid switching between motor plans [17]. Phase switches have been observed when trials alternate between a series of reaches with a misaligned cursor and subsequent no-visual feedback trials [18]. While this transition has often been attributed to the removal of an explicit strategy, recent results [19] suggest that it cannot be fully explained in this way. Our current work shows that this behavior can instead be captured within a computational framework.

A particular challenge in modeling visuomotor adaptation lies in capturing the behavioral consequences of varying participant instructions and the nonlinear saturation observed in paradigms such as error clamp experiments. To address this, we designed a visuomotor reaching task that introduces a novel manipulation: a hand angle-gain parameter, denoted by *α*, which scales the angular displacement of the hand relative to the target, independently of the standard visuomotor rotation applied to the visual cursor. See Fig 5.

This gain-based manipulation allows us to implement error clamp conditions in a graded, parametric manner. Specifically, *α* = 0 corresponds to a full error clamp, while *α* = 1 corresponds to standard adaptation conditions. By systematically varying *α* between 0 and 1 — assigning a fixed gain per experiment — we observe the progressive emergence of saturation effects. This graded approach offers a principled way to study and model nonlinear learning behavior. To our knowledge, this experimental method has not been previously employed; prior work has focused primarily on error augmentation paradigms [20, 21]. Our findings suggest that the error clamp should not be treated as a singular or exceptional case, but rather as an integral part of the behavioral response continuum — one that any comprehensive model of visuomotor adaptation must account for.

## 2 Materials and Methods

### Participants

Experiments 1 and 2 involved 29 participants (males = 15, females = 14, age range = 18 −32, age (mean ± standard deviation) = 21.69 ± 3.41 years). Experiments 3 and 4 involved 51 participants (males = 20, females = 31, age range = 18 −38, age (mean ± standard deviation) = 21.7 ± 4.73 years). All the participants self-reported to be right-handed or left-handed, and having normal or corrected-to-normal vision. Also, prior to the data collection, all participants signed and were informed with a written consent form. This study was approved by York University’s Human Participants Review Committee and all procedures were performed in accordance with institutional and international guidelines.

### Experimental Setup

The setup is illustrated in Fig 4 and consisted of a downward-facing LCD screen (60 Hz, 20 in, 1680 × 1050, Dell E2009Wt), a tablet and stylus (Wacom Intuos Pro PTH860) to record the hand position, and an upward facing mirror placed in between the tablet and screen. In-house software was created to design and conduct visuomotor reach experiments (https://github.com/thartbm/PyVMEC2). Visual stimuli were presented via a downward-facing LCD screen positioned above an upward-facing mirror, such that the reflected image appeared aligned with the horizontal plane as the participants’ hand. Participants were seated in a height-adjustable chair, allowing them to comfortably execute out-and-back reaching movements on the tablet by using a stylus held in their dominant hand. A black cloth was placed over the shoulder to occlude vision of the arm.

### Visuomotor Reaching Task

Participants made outward reaches toward a target by smoothly gliding the stylus on the tablet starting from a home position, as shown in Fig 4. Each trial began at a central home position, displayed as an open gray circle (radius 0.5 cm) on the screen. The reach target, also an open gray circle of the same size, appeared 8 cm from the home position, generally in the forward direction. The target appeared when the stylus entered the home position. Participants were instructed to move toward the target at a comfortable pace — neither too fast nor too slow — and to avoid resting their wrist or elbow on the tablet or surrounding surface during the reach.

Two types of trials were included, based on task instructions. In **Learn** trials, participants were instructed to move the *cursor* to the target, promoting adaptation to visuomotor errors. In **Ignore** trials, they were instructed to move their *unseen hand* to the target location, regardless of cursor position, thus minimizing reliance on visual feedback. Both trial types were performed under continuous cursor feedback. The first two experiments also included **No Cursor** trials in which a cursor was not presented and participants were instructed to move the unseen hand directly to the target. The second two experiments included **Washout** trials with veridical cursor feedback and instructions to move the cursor to the target.

In trials with visual cursor feedback, a black cursor (radius 0.5 cm) continuously represented the stylus position, allowing participants to track their unseen hand. In no cursor trials, the cursor was not visible during the reach. To encourage consistent movement speeds, auditory feedback was provided based on the movement duration. Outward movements lasting between 0.2s and 0.4s triggered a pleasant tone, while movements that were too fast, too slow, or initiated before target onset triggered an unpleasant tone. At the end of each trial, participants were instructed to return their hand to the home position to initiate the next trial. As the hand is not visible, a stencil was used to guide the hand back to the start location reliably. The stencil outline is depicted as a cone shaped region in Fig 4. The tip of this cone coincides with the home position, such that participants can return to the home position without any visual information.

### Hand Angle-Gain

Next we explain the rule used to determine the cursor direction for Learn and Ignore trials. Let *d* denote the perturbation (rotation in degrees), *r*(*k*) the direction of a target on the *k*-th trial, and *x*(*k*) the hand angle on the *k*-th trial. We introduce a parameter *α* called the *hand angle-gain* that takes one of four values in the first two experiments: *α* ∈ {0.4, 0.6, 0.8, 1.0}. The cursor was placed at the following angle

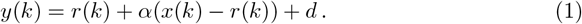

This rule ensured that the cursor was displayed at an angle offset by the rotation *d*, while the hand’s contribution to that offset was scaled by *α*. When *α* = 1.0, the cursor directly reflects the rotated hand angle — standard visuomotor rotation. As *α* decreases toward 0, the cursor movement is increasingly clamped to the target direction plus the perturbation, effectively simulating error clamp conditions.

Fig 5 illustrates the behavior of the cursor direction for a target positioned straight ahead at *r* = 90° and a perturbation *d* = 15°. In the left panel, the participant initially aims for *x*(*k*) = 90°, causing the cursor to appear at *y*(*k*) = *r* + *d* = 105°. In the middle panel, the participant compensates for the initial error by adjusting their reach in the direction opposite to the perturbation. According to (1), the hand angle is scaled by parameter *α*, which moves the red arrow closer to the target. This adjusted hand angle is then rotated by *d* = 15° to produce the cursor angle, shown as graded blue arrows, each corresponding to a different *α* value. As *α* decreases, the hand angle is progressively shifted towards the target angle, and the cursor angle moves closer to the original cursor angle shown in the left panel. When *α* = 0, the hand angle is fully suppressed (multiplied by zero) and rotated by 15°, resulting in a constant cursor angle at *y* = 105°, a condition known as the error clamp [22]. Conversely, when *α* = 1, the participant experiences a standard learning trial without any scaling of the hand movement. The right panel of Fig 5 illustrates the hand angle required to fully compensate for the perturbation, effectively eliminating the observed error and aligning the cursor with the target.

### Experimental Structure

The experimental study included four experiments. In Experiments 1 and 2, each participant completed the visuomotor reaching task across two experimental sessions, separated by a minimum of three days. The experimental structure was identical across sessions to assess within-participant consistency and to ensure adequate data for analysis. In each session, participants performed two distinct experiments (1 and 2). The four cycle structure of Experiments 1-2 primarily allowed us to collect data for four values of *α* in a single session. From the point of the view of the model, there is no relationship between cycles that is being probed with this experimental structure. Indeed, we alternate the sign of the perturbation between cycles to avoid the possibility of carry-over from one cycle to the next. In Experiments 3 and 4, participants completed only one type of experiment in one session only.

As illustrated in Fig 6, Experiments 1 and 2 were organized into four sequential cycles, indicated by vertical dashed lines. Each cycle consisted of three phases: a baseline phase, a rotation phase, and a no cursor phase. A brief one-minute break was provided between cycles in both experiments. During this time, on-screen instructions were presented to remind participants of the upcoming task requirements. Experiments 3 and 4 consisted of only one cycle with three phases.

- **Experiment 1** (**Ignore-N Experiment**) consisted of four cycles, each with three trial phases (Fig 6). Each cycle began with a baseline phase of 40 trials in which the cursor accurately represented the stylus position (i.e., veridical feedback, no rotation). This was followed by a phase of **Ignore** trials, in which participants were instructed to move their *unseen hand* to the target while a visuomotor rotation (perturbation) was applied to the cursor. In addition to the perturbation, an angular gain factor — referred to as the *hand angle-gain α* — was applied to the hand trajectory to manipulate the relationship between the hand angle and the resulting cursor angle. Further details about the implementation and rationale for *α* are provided below. Each cycle concluded with a phase of 40 **No Cursor** trials, during which the cursor was not visible throughout the reach. Targets in Experiment 1 were presented at one of five angular locations: 80°, 85°, 90°, 95°, or 100°, relative to the center. The Ignore phase of Experiment 1 introduced both a ± 15° visuomotor rotation as well as a hand angle-gain *α*. To minimize carryover effects in this within-subject design, the sign of the rotation alternated across cycles (e.g., +15°, −15°, +15°, −15°). Additionally, across the four cycles, *α* took values of 1.0, 0.8, 0.6, or 0.4, allowing a progressive attenuation of the effect of the hand movement on the cursor placement — approaching an error clamp as *α* is decreased (see Fig 5). The hand angle-gain *α* was always set to 1.0 during the baseline phase, and it is not relevant in the no cursor phase. The number of Ignore trials within a cycle depended on *α*. For *α* = 1.0, 0.8, or 0.6, the Ignore phase contained 100 trials. For *α* = 0.4, the number of Ignore trials was increased to 160 to allow for sufficient data under near-clamp conditions.
- **Experiment 2** (**Learn-N Experiment**) followed a similar structure as Experiment 1, with each cycle beginning with 40 baseline trials and concluding with 40 no cursor trials. The central phase of each cycle consisted of 100 **Learn** trials, in which participants were instructed to move the *cursor* to the target. During the Learn phase, both a rotation and a hand angle-gain were applied to the cursor, following the same pattern as for Experiment 1. Targets in Experiment 2 were presented at one of five angular locations: 80°, 85°, 90°, 95°, or 100°, relative to the center.
- **Experiment 3** (**Ignore-W Experiment**) included only one cycle with three phases. The experiment began with 40 baseline trials and concluding with 40 **Washout** trials with veridical cursor feedback. The central phase of this experiment consisted of 120 Ignore trials. During the Ignore trials, the perturbation was either 25° CW or CCW, and the value of *α* was one of {0.5, 0.75, 1.0}. Five targets were placed at every 5° from 140° to 160° for a CCW perturbation; while five targets were placed at every 5° from 20° to 40° for a CW perturbation. Each participant experienced either a CW or CCW rotation, but not both, and only one value of *α* within the session.
- **Experiment 4** (**Learn-W Experiment**) had the same structure as Experiment 3, but participants performed Learn trials instead of Ignore trials.

### Data Analysis

The performance of each participant was evaluated by measuring the angular reach deviation for each trial. The angular deviation was computed as the angle between a line through the target and a line through the point of maximum velocity of the hand. Baseline biases for each participant were calculated by using the average angular reach deviation over the last 20 trials of the baseline phase. The angular reach deviation of each trial was then corrected based on each participant’s baseline biases.

Finally, the mean reach deviation was computed over all participants using the bias-corrected data. All raw and summary data are publicly available: https://osf.io/gjb5r/.

In the experiments, the target was placed at one of five randomized locations; for instance, between 80° and 100° in Experiments 1 and 2. However, for the simulations, we normalize the data to treat the target as if it were always at 90°, representing a single direction within the adaptation field. For the experimental data from other studies, hand angles were averaged over targets and over participants. A bias correction using the baseline phase of the averaged participant data was only applied to the data from [23].

## 3 Model Components

The paper presents a nonlinear state-space model of the visuomotor reach experiment called the **Disturbance Observer** (DO) Model, a metonym for the most significant component of the model, a disturbance observer. This section introduces the model components, including intuition for each computational module and design principles that inform the approach. We also present a preliminary visuomotor model to provide insight on the model’s nominal operation. The section concludes with a summary of the modeling assumptions.

We begin by directing the reader to several overview materials to obtain a bird’s eye view of the model. Table 1 summarizes the signals in the model, their meaning and origin, whether evolving in the physical world (including the body) or in the brain. Table 2 summarizes the model parameters, including their role in the model. Figures 1-3 show three modalities that the model is intended to capture: Learn trials when a cursor is visible, Ignore trials when a cursor is visible, and No Cursor trials when moving the hand to the target. This section of the paper will focus on Figure 1. All angular quantities in the DO model are in units of degrees, with 90° representing the straight ahead direction - defined as a horizontal line through the participant’s sagittal plane while seated.

**Table 1.** Classifications of Signals. Signals associated with the plant model arise from four sources: the body, specifically the hand; the environment, in the form of the perturbation; the cursor angle, interpreted as the output of the plant; and the measurement, the visual error observed by the subject. Signals associated with the three control modules all arise in the brain.

|  | Signal | Source | Role in Model |
| --- | --- | --- | --- |
| Plant Model | $x(k)$ | body | hand angle |
| | $d(k)$ | environment | perturbation |
| | $y(k)$ | output | cursor angle |
| | $e(k)$ | measurement | visual error |
| Error Feedback | $u_s(k)$ | brain | single-trial learning |
| Disturbance observer | $w_0(k)$ | brain | disturbance observer state |
| | $\hat{w}(k)$ | brain | estimate of perturbation |
| | $u_{im}(k)$ | brain | disturbance observer motor command |
| Feedforward system | $x_f(k)$ | brain | feedforward fast state |
| | $r_f(k)$ | brain | feedforward slow state |
| | $u_f(k)$ | brain | feedforward motor command |

**Table 2.** Nominal model parameters.

| Module | Parameter | Nominal Values | Role in Model |
| --- | --- | --- | --- |
| Error Feedback | $K$ | 0.25 | error sensitivity in single-trial learning |
| Disturbance observer | $F$ | 0.7 – 0.9 | learning rate of disturbance observer |
| | $G$ | $1 - F$ | requirement of internal model principle |
| | $\psi_o$ | 0.9 – 1.0 | proportion of learning at asymptote |
| | $b_w$ | 0.001 | saturation strength |
| Feedforward system | $A_f$ | 0 – 0.9 | learning rate of feedforward system |
| | $A_{fn}$ | 0.95 | forgetting rate during no cursor trials |
| | $L_o$ | 1.1 | error sensitivity to $u_{im}$ |
| | $b_f$ | 0.01 | saturation strength |
| | $L_f$ | 0.0001 | learning transfer rate |

**Fig 2.**
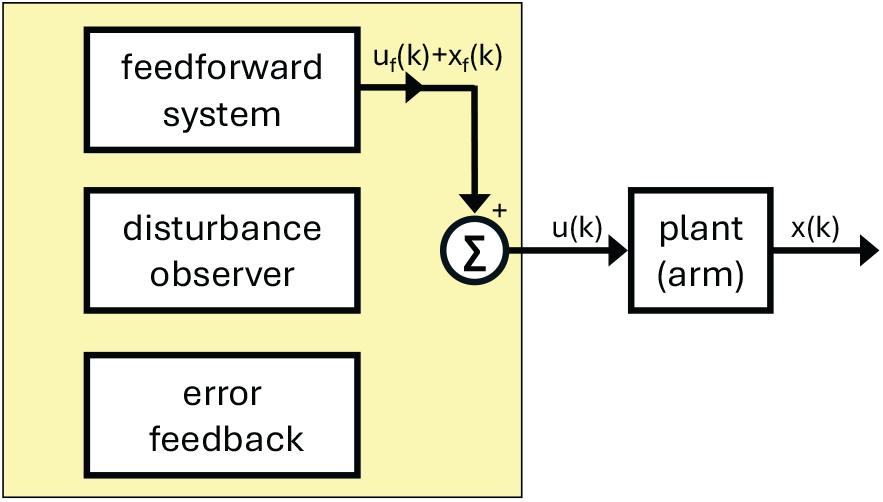
Block diagram of the DO model during No Cursor trials.

**Fig 3.**
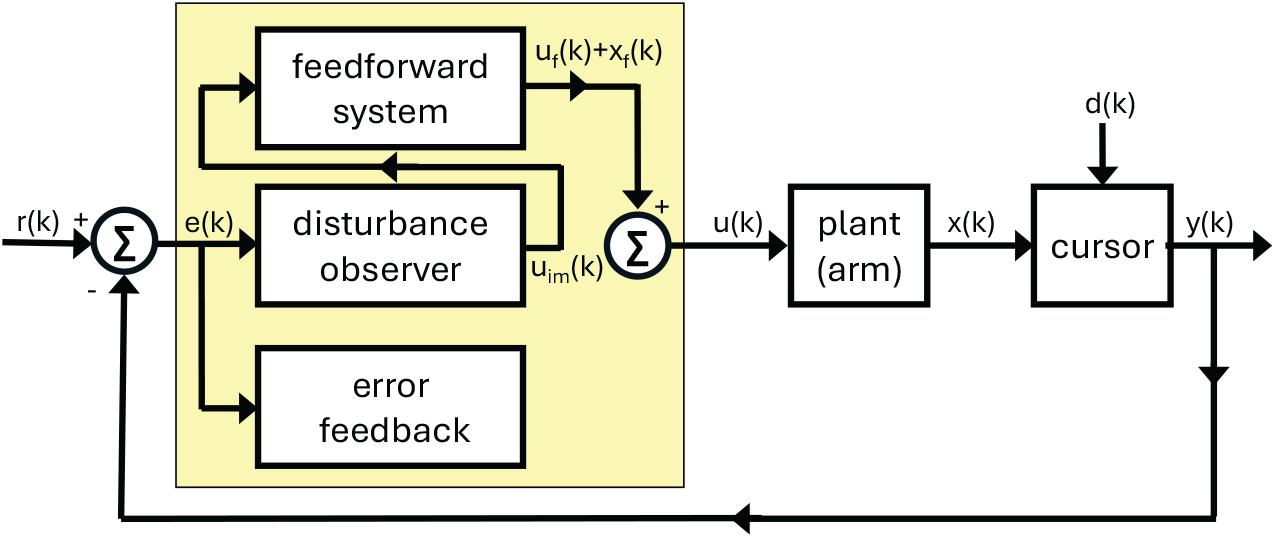
Block diagram of the DO model during Ignore trials.

**Fig 4.**
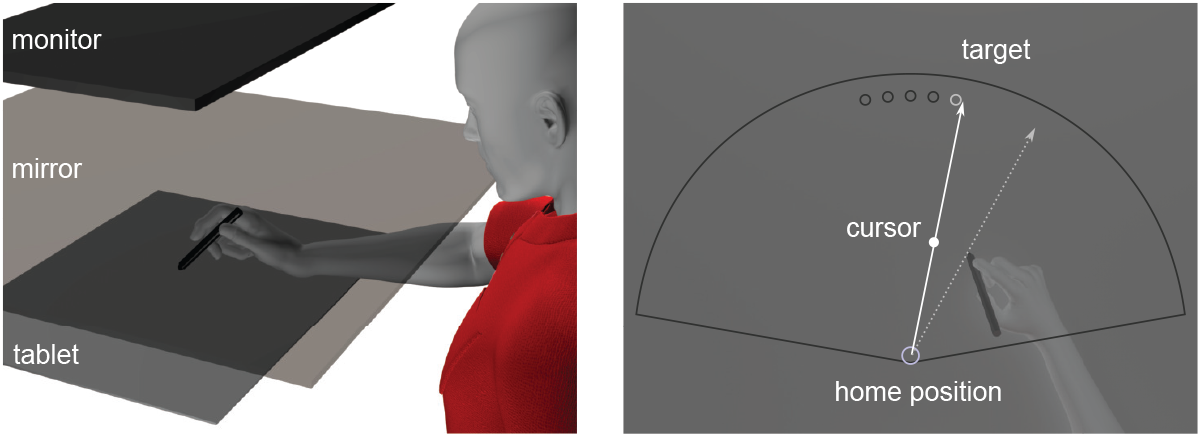
(a) Experimental apparatus for presenting a visuomotor reaching task. (b) Representation of a trial with visual feedback.

**Fig 5.**
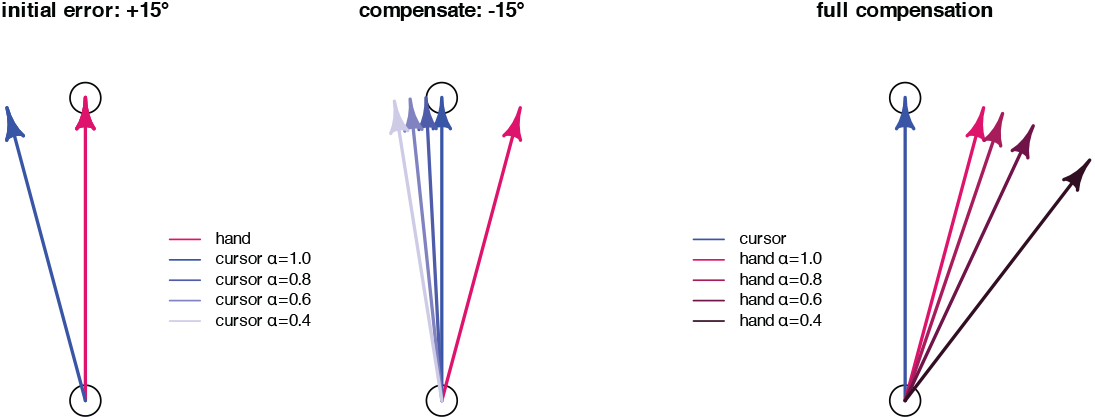
Effect of the hand angle-gain (*α*) on the cursor direction. The left figure shows the initial observed visual error due to a perturbation of +15°. The middle figure shows the compensation of the hand angle by − 15° to overcome the initially observed visual error of magnitude +15° with *α* = 1.0. The shaded blue arrows show the effect on the cursor direction for *α* ∈ {0.8, 0.6, 0.4} with lighter blue for smaller *α*. We see that as *α* is decreased, the cursor direction progressively approaches a constant value of 115°. The right figure depicts the required compensation of the hand angle to place the cursor on the target for *α* ∈ {1.0, 0.8, 0.6, 0.4}.

**Fig 6.**
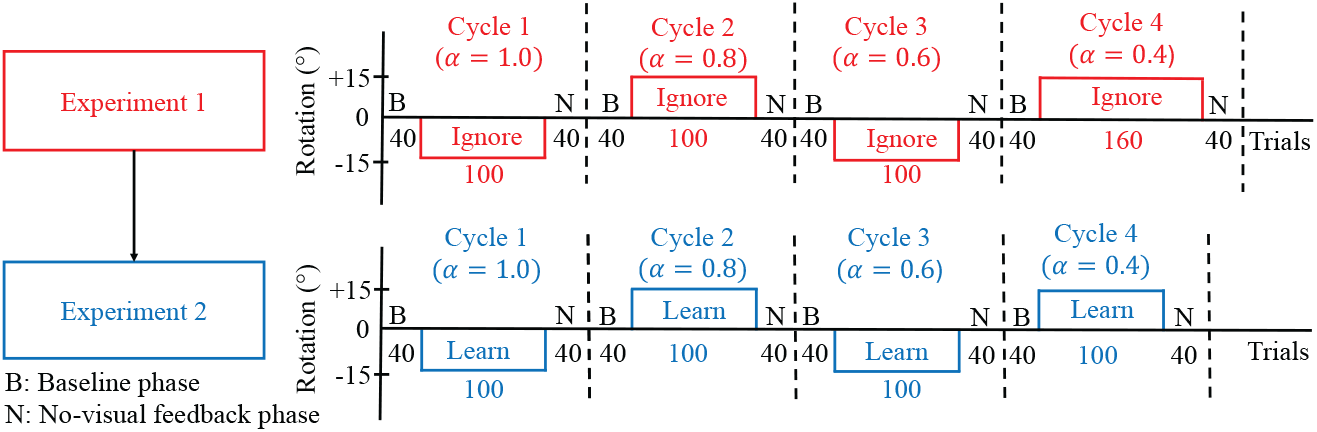
Trial structure for Experiments 1 and 2, with Experiment 1 performed first. In Experiment 1 (Ignore-N) participants were instructed to move their invisible hand to the target while ignoring the cursor. The experiment was conducted in four cycles, denoted cycle 1 to cycle 4. Each cycle began with a baseline phase (B) of 40 trials with no rotation and a hand angle-gain (*α*) = 1.0, followed by a rotation phase of Ignore trials with ± 15° rotation. Each cycle ended with a no cursor phase (N) of 40 trials with no rotation and *α* = 1.0. In the Ignore trials of each cycle, participants experienced one of four different values of *α*: 1.0, 0.8, 0.6, and 0.4. In Experiment 2 (Learn-N) participants were instructed to move the cursor to the target. The experiment was also conducted in four cycles. Each cycle began with a baseline phase (B) of 40 trials with no rotation and *α* = 1.0, followed by a rotation phase of Learn trials with ± 15° rotation. Each cycle again ended with a no cursor phase (N) of 40 trials with no rotation and *α* = 1.0. In the Learn trials of each cycle, participants experienced one of four different values of *α*: 1.0, 0.8, 0.6, and 0.4. There was a one-minute break in between each cycle of these experiments, and during that time, particular instructions were displayed on the screen for the upcoming trials.

Considering Figure 1, we see that the overall model structure is that of a classical unity feedback loop, the most ubiquitous control architecture of control theory. Moving around this control loop, we start with the plant, the mechanism being controlled, which in the considered visuomotor experiments of the paper is the arm. The hand angle *x*(*k*) on trial *k* is combined with an exogenous perturbation *d*(*k*) according to a formula, determined by the experimenter, to place a cursor on a digital pad, with a cursor angle *y*(*k*). The cursor angle can be interpreted abstractly as the output of the plant. The visual error which is recorded in the brain is modeled as the difference between a target angle *r*(*k*) and a cursor angle *y*(*k*). The controller (depicted as a yellow box) represents salient computations in the brain to move the arm, focusing on the error-driven computations required in visuomotor adaptation. Importantly, the controller receives only a single measurement to perform its computations, which is the visual error *e*(*k*) on trial *k*. The controller consists of three sub-modules, which have been organized by timescales, with the fastest timescale at the bottom and the slowest timescale at the top. The output of the controller is the motor command *u*(*k*), which determines the hand angle for the next reach. We now describe how to model each component in this control architecture.

### 3.1 Model of the Plant

The plant is the mechanism to be controlled, which in visuomotor reach experiments is the arm. The primary modeling task is to model the transformation from control inputs to plant states and outputs. The modeler has a choice to use a relatively detailed mechanical model of the arm or to use a more abstract discrete time model of the input-output behavior of the arm over successive trials of the experiment.

A reasonably detailed mechanical model of arm motion is given by:

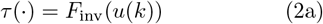

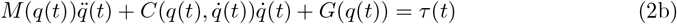

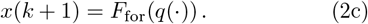

Equation (2b) is an Euler-Lagrange model of the arm, widely used in robotics and biomedical applications to capture the physics of arm movement over time as a function of torque inputs. The vector *q*(*t*) ∈ ℝ^*n*^ represents the continuous-time joint variables, including the rotational degrees of freedom of the shoulder, a joint variable for the elbow, and the rotational degrees of freedom of the wrist. The continuous time open-loop torques *τ* (*t*) ∈ ℝ^*n*^ represent the aggregate effect of the muscle groups that move each joint. These torques are computed in open-loop in the sense that it is assumed that the subject moves fast enough on each arm reach so that there is no mid-course correction of these torques.

In the DO model, the motor command *u*(*k*) represents the brain’s commanded hand angle computed at the end of trial *k* for the next arm reach at trial *k* + 1. The interpretation of (2a) is to resolve the passage from a discrete-time motor command based on data up to and including trial *k* to the continuous time torques applied at the joints to achieve that command on the next trial. The full torque trajectory as a function of time for one trial is represented by *τ* (·). The computation in (2a) involves a linear-in-parameters internal model of the arm dynamics as well as the inverse kinematics of the arm; both well-known computations [24].

Equation (2c) models the passage from continuous-time evolution of arm movement on each trial to a discrete-time state. The full trajectory for *q*(*t*) over a single arm reach on trial *k* + 1 is represented as *q*(·). This trajectory is converted to a 2D motion of the hand on the digital pad using the forward kinematics of the arm. Finally, the model computes the discrete state *x*(*k* + 1), the hand angle on trial *k* + 1 as the angle of the hand with respect to a line through the target at the time of maximum speed of the continuous hand movement. All these computations are embedded in the formula (2c).

We notice that equations (2b) and (2c) model the physics of the arm and of the experiment, whereas (2a) models a computation performed in the brain. For researchers who are specifically interested to study computations associated with internal models of the plant, (2a) is the key equation that must be more fully elaborated in the model. On the other hand, for subjects who have not recently been injured, the brain computation in (2a) may be assumed to incorporate plant parameter estimates that are closely matched to the true plant parameters in (2b). If we assume that plant and internal model parameters are exactly matched, then mathematically the transformation from *u*(*k*) to *x*(*k* + 1) is the identity function. In other words, for healthy subjects the transduction from commanded hand angle *u*(*k*) to actual hand angle *x*(*k* + 1) may be idealized as being close to perfect. This leads to an alternative trivial plant model

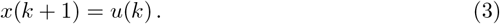

This trivial plant model assumes that the computation in (2a) is protected from short-term environmental factors, so that (2a) remains an accurate inverse computation throughout the experiment. This important assumption will be revisited at the end of this section, as we review all modeling assumptions.

We strongly recommend that modelers utilize the trivial plant model (3) in cases when they are not explicitly concerned with detailed plant dynamics or with internal models of the plant. This focuses the model on the error-driven dynamic process that regulates the visual error to zero over successive Learn trials, which we regard as the core computation in visuomotor adaptation.

### 3.2 Model of Cursor Placement and Visual Error

The next modeling step is to model the placement of the cursor on each trial in the experiment and to model the visual error that the subject perceives. Let *r*(*k*) denote the angular position of the target, *d*(*k*) denotes the angular perturbation, and *y*(*k*) denotes the cursor angle, all on the *k*-th trial. The standard formula for placement of the cursor is *y*(*k*) = *x*(*k*) + *d*(*k*). That is, the true hand angle *x*(*k*) is rotated by a perturbation *d*(*k*). Notice that this formula does not represent a modeling choice, but rather captures the precise formula used in the experimental runtime software. The formula announces that we are not studying force-field experiments, in which case the perturbation would instead arise as an additive term in (2a).

We noted in Section 2 that Experiments 1-4 in this paper use a more involved cursor placement rule given by (1). The scale factor *α* in (1) introduces a *hand angle-gain* relative to the target. We restrict *α* to the range *α* ∈ [0, 1] in this paper. Its behavior has been described in Fig 5, which illustrates the impact of the hand angle-gain on the cursor’s direction. First, consider the case when *d*(*k*) = 0. We can rewrite (13b) as:

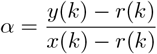

Thus, *α* determines the ratio of the cursor angle to the hand angle, both relative to the target angle. To compensate for this reduced angular gain, participants must produce a larger hand deviation - an effect illustrated in the right panel of Fig 5, with smaller *α* depicted by darker red arrows.

Returning to (1), we have chosen not to scale *d*(*k*) by *α* so that we can independently manipulate the hand angle-gain and the perturbation size in our experiments. An additional motivation for introducing *α* comes from considering its limiting values of *α* = 0 or *α* = 1. Suppose *α* = 0, and *r*(*k*) ≡ *r* and *d*(*k*) ≡ *d*, both constants. Then the cursor angle is

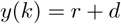

which corresponds to an error clamp condition in which the cursor angle is independent of the hand angle. When *α* = 1, the cursor angle is

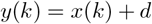

which corresponds to a standard visuomotor rotation paradigm. By decreasing *α* from 1 to 0, it becomes possible to gradually introduce an error clamp condition - referred to as a *graded error clamp*. The error clamp is known to elicit a saturated adaptive response [22, 25]; thus, varying *α* allows us to observe the emergence of nonlinear effects incrementally, across a series of experiments with different *α* values.

The standard model of the visual error, ubiquitous in the eye movement literature [26], is

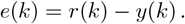

This equation captures that the retina records the cursor angle relative to the target angle, with a scale factor that has not been included in the model.

### 3.3 Model of Feedback Controllers

The heart of the DO model is the model of the controller, which operates on three timescales. As seen in Figure 1, the output of the controller is the overall motor command *u*(*k*), the sum of all active control modules for a particular trial type (Learn, Ignore, No Cursor). We see that for Learn trials, all control modules are active. Formally speaking, the model has two feedback control modules, described in this subsection. The third control module, the feedforward system, is formally not a feedback controller because it does not directly receive the exogenous error measurement *e*(*k*).

#### 3.3.1 Error Feedback

The fastest control module at the bottom of Figure 1 is an error feedback controller, modeled as:

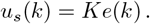

The name “error feedback” refers to the mathematical structure of this formula; namely a gain parameter *K* multiplies an error *e*(*k*). The error is fed back to the input side of the plant; the term “feedback” may have other connotations in the neuroscience literature which do not apply here. The notation *u*_*s*_(*k*) is intended to recall the fact that, in control-theoretic terms, this component of the motor command is for stabilization of the closed-loop system. In the experimental setting, it captures how participants adjust their movement following a visual error on the previous trial. As such, one can think of *u*_*s*_(*k*) as modeling *single-trial learning* [27–29]. The parameter *K* is interpreted as the participant’s *error sensitivity* [30]. The effect of *u*_*s*_(*k*) is observed only in the early phase of learning, when errors are large.

#### 3.3.2 Disturbance Observer

A disturbance observer is a type of internal model of persistent exogenous disturbance signals [31]. This type of internal model contrasts with internal models of the plant, as seen in (2a), which are currently more common in the neuroscience literature. An internal model of an environmental signal only confers benefits to a control system if that environmental signal is predictable. Predictable disturbance signals (in continuous or discrete-time) include step signals (constant signals), ramp signals, sinusoidal signals, multisinusoids, and linear combinations of these. This class of signals is sufficiently rich to capture most (non-random) persistent disturbances in robotics and motor control applications. In Learn trials of the visuomotor reach experiment, the perturbation is most commonly a constant disturbance signal that enters into the visual error. Therefore, there is justification to use an internal model approach to model how the brain estimates this disturbance; in fact, there is compelling evidence that the visuomotor system is unable to estimate even a mildly more complex disturbance such as a sinusoid [32].

There are many internal model designs available in the control literature for dealing with exogenous disturbances affecting a control loop. We have chosen to work with a design from [31] due to its simplicity. We will say more about alternative design approaches in Section 6.2. We must emphasize that the disturbance observer proposed in this paper is only able to estimate constant perturbations, corresponding to the visuomotor experiments we considered in which learning takes place under this condition. The disturbance observer model is given as:

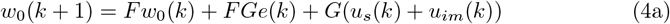

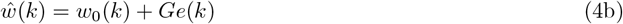

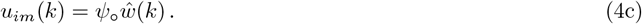

A single (scalar) state *w*_0_(*k*) ∈ ℝ is associated with the disturbance observer. Its update in (4a) relies on an efference copy of two components of the motor command as well as a measurement of the visual error. All three signals are assumed to be available in the brain. The output of the disturbance observer is the signal *ŵ* (*k*) ∈ ℝ, which is formed as a linear combination of the error *e*(*k*) and the state *w*_0_(*k*) in (4b). The overall output of this control module, as seen in Figure 1, is the signal *u*_*im*_(*k*), the component of the motor command whose specific function is to cancel the perturbation appearing in the visual error.

The disturbance observer has two free parameters *F* and *ψ*_o_. The parameter *F* sets the learning rate of the disturbance observer, and its effect is seen only in the transient phase of learning. The parameter *ψ*_o_ is a scale factor that modulates the amount of learning at asymptote. If *ψ*_o_ = 1, the net effect of *u*_*im*_(*k*) is to drive the visual error to zero over successive Learn trials. The parameter *G* is not a free parameter, and it is defined as

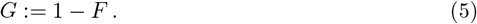

It’s role is to ensure the internal model principle is satisfied, as explained below. The full motor command consists of three components:

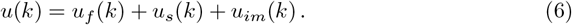

The first component corresponds to an intact open-loop feedforward system that corresponds to the transduction from the visual perception of a target in the visual field to a high-level motor plan to move the hand directly to the target. This component is not an error driven computation, but rather captures a precalibrated visual mapping between what is seen in the world and the selection of a high level motor plan. In the healthy subject, this precalibrated visual mapping stored in the brain will be close to perfect. As such, we model it as being ideal:

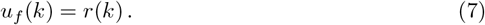

This equation says that the high-level plan that the brain has a priori calibrated provides a correct correspondence between the retinotopic field and physical space. Returning to (6), we see that *u*(*k*) consists of a nominal high level plan *u*_*f*_ (*k*) which is pre-computed in the brain and is available for use by the healthy subject, whereas the components *u*_*s*_(*k*) and *u*_*im*_(*k*) provide two error correction mechanisms to the nominal plan: a fast correction *u*_*s*_(*k*) and and slower correction *u*_*im*_(*k*).

To understand how the disturbance observer works, we consider its computations over a sequence of Learn trials with the cursor visible, with the target at a fixed angle *r*(*k*) ≡ *r*, and the perturbation a constant signal *d*(*k*) ≡ *d*. Referring to (1), we also suppose *α* = 1. Under these assumptions, the visual error is

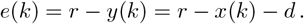

Consider the update of the signal *ŵ*(*k*) on the next trial:

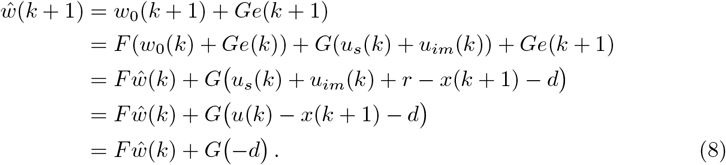

This calculation shows that the hand movement *x*(*k*) and the motor command *u*(*k*) have been exactly canceled through updates performed by the disturbance observer. This cancellation isolates the only unexpected component in the visual error, namely the perturbation *d*. Specifically, the formula (8) with *F* ∈ (0, 1) is a stable filter, with a rate of convergence determined by *F*. This filter admits a well-defined steady state. Since *G* = 1 − *F*, the steady state value of *ŵ*(*k*) is

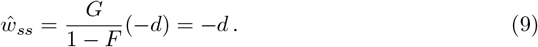

In summary, under nominal operational conditions, the disturbance estimate *ŵ*(*k*) converges to the negative of the perturbation.

Importantly, when the perturbation *d*(*k*) varies over time or includes random fluctuations, then this disturbance observer will struggle to form a reliable estimate of the perturbation - consistent with empirical findings that environmental consistency facilitates learning [32–35]. Secondly, if a subject had sustained a recent injury, then correspondingly in the model, the cancellation of *x*(*k* + 1) and *u*(*k*) in the update of *ŵ*(*k* + 1) would not be perfect. The performance of the disturbance observer would be seriously degraded in this case.

An important feature of the disturbance observer is its recurrent architecture, as seen by the fact that the output *u*_*im*_(*k*) of this control module becomes an input to the disturbance observer in (4a). This recurrent architecture is crucial for the satisfaction of the internal model principle. To see this, substitute *u*_*im*_(*k*) into (4a) to get:

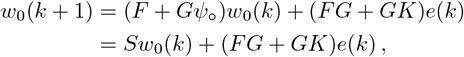

where *S* := (*F* + *Gψ*_o_). This update of the state *w*_0_(*k*) has the form of the classical internal model of E.J. Davison [36, Eq. 10], here for discrete-time systems. The internal model principle is satisfied if all neutrally stable poles of the *z*-transform of the signal *d*(*k*) are eigenvalues of *S*. Since *d*(*k*) is assumed to be a constant signal, it has a single neutrally stable pole at *z* = 1 in the complex plane. Therefore, we require that *S* = 1. If we assume a value of *ψ*_o_ = 1 to model complete learning, then we require *G* = 1 − *F*, which is the value we selected in (5).

### 3.4 Preliminary Visuomotor Model

We have proposed two feedback controllers and one a priori tuned feedforward system to represent computations in the brain to perform visuomotor adaptation. One can now build a preliminary visuomotor model that is specialized to handle Learn or Washout trials (the cursor is always visible and the instructions are to move the cursor to the target). We consider this preliminary model:

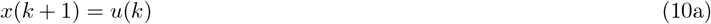

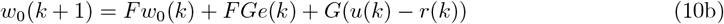

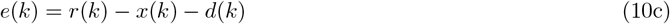

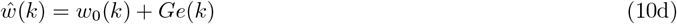

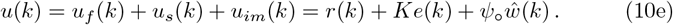

This preliminary model has three free parameters: *K* for fast learning, *F* for slow learning, and *ψ*_o_ for the proportion of learning at asymptote. We have deduced from (8) that under the assumptions *r*(*k*) ≡ *r* and *d*(*k*) ≡ *d*, the disturbance observer forms a stable filter computation that estimates the perturbation. Therefore, a reduced model with fewer signals is:

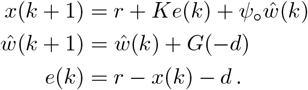

This model may be further simplified by writing an *error model* for *e*(*k*); namely a model that describes the dynamic evolution of the error:

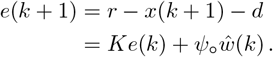

This error model replaces the dynamic model of the hand angle (the choice of states is not unique), so we obtain a simplified closed-loop model

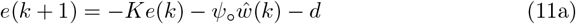

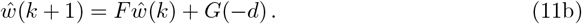

This closed-loop model is guaranteed to be stable if we select *F* ∈ (0, 1) and *K* ∈ (0, 1). In turn this means that a steady-state exists. First, we have *ŵ*_*ss*_ = −*d*, based on (9). Next, compute the steady-state error and hand angle:

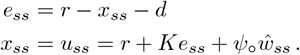

Substituting *e*_*ss*_ and *ŵ*_*ss*_ = −*d* into the formula for *x*_*ss*_ and solving for *x*_*ss*_, we obtain

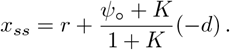

If *ψ*_o_ = 1 to model complete learning, then *x*_*ss*_ = *r* − *d* and *e*_*ss*_ = 0. This analysis explains how the preliminary model learns the full perturbation in order that the steady-state error is driven to zero. The essential requirement to achieve this feat is the internal model principle of control theory.

Stepping back, we started with a preliminary visuomotor model (10) with a trivial model of the plant (representing perfect execution of the high level plan), two feedback controllers modeling error-driven computations in the brain, and a perfectly calibrated feedforward system (not including biases). We performed two coordinate transformations to arrive at a simpler closed-loop model (11). A key observation is that (11) is simply a variant of the two-rate model. The only difference is that we have utilized states that have a physical meaning (again, the choice of states is not unique). The fast state is *e*(*k*), and it evolves in the world. The slow state is *ŵ*(*k*), and it evolves in the brain. Further illuminating comparisons with the two-rate model are found in Section 6.3.

### 3.5 Model of Feedforward System

We have proposed a preliminary visuomotor model (10) whose underpinning is the internal model principle of control theory. This model is functionally equivalent to a two-rate model, as seen by performing two coordinate transformations. Unsurprizingly, (10) is not an advancement beyond the two-rate model in terms of expressivity, though it does clarify the nature of the computations in the brain. We would like to develop a model with greater expressivity, and the key behavior that is missing is the capability to move the hand to the target when there is no visual error: No Cursor trials.

The idea that No Cursor trials would be driven directly by the disturbance observer is improbable. The disturbance observer only learns when a visual error is present, and experimental evidence (that perturbations cannot be unlearned without washout trials) suggests it remains in a quiescent state until further information is provided to it. Moreover, if the goal during No Cursor trials is to move the hand the target, then disturbance observer signals contain erroneous information about a perturbation than no longer exists. We are lead to the commonsense deduction that a new computational module, capable to operate in open-loop (without an error signal), is needed to characterize the high-level computations in the brain for No Cursor trials.

The feedforward system is a motor system for reaching the hand along a specific direction (or to a specific location) without the benefit of error feedback, for instance when we close our eyes, when the hand is occluded, when we reach behind the back, and so forth. Like any motor system, the feedforward system requires ongoing calibration - it cannot operate only in open-loop. The present model conceives this calibration to arise through training by the disturbance observer. Two points must be considered for modeling the feedforward system: (i) the model must capture that subjects already have a pre-calibrated feedforward system when they initiate a visuomotor experiment; (ii) learning takes place in the feedforward system based on the accrued experience of the disturbance observer.

The first point has already been discussed when we introduced the equation (7), a modeling assumption which can be related to assumptions about the plant. However, choosing the plant model to be *x*(*k* + 1) = *u*(*k*) regards the motor side of the visuomotor system. It means the motor plan to move in a direction *u*(*k*) is correctly converted to torques that move the hand. Instead, *u*_*f*_ (*k*) = *r*(*k*) is a statement about the sensory side of the visuomotor system. It means that information recorded on the retina is correctly transformed into a motor plan.

The second point involves modeling how the feedforward system learns from the disturbance observer. We propose the following model:

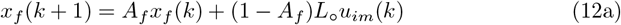

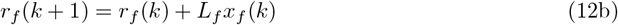

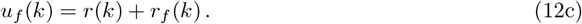

We see that there are two states associated with the feedforward system: *x*_*f*_ (*k*) and *r*_*f*_ (*k*). Learning initially takes place in *x*_*f*_ (*k*), driven by the output of the disturbance observer *u*_*im*_(*k*). The rate of learning is determined by parameter *A*_*f*_, while *L*_o_ is the sensitivity to *u*_*im*_(*k*). Learning in *x*_*f*_ (*k*) is slowly transferred to the second state *r*_*f*_ (*k*) where *L*_*f*_ ≪ 1 is a small parameter representing the transfer rate. Finally, the feedforward motor command *u*_*f*_ (*k*) is slowly updated with the correction *r*_*f*_ (*k*) to the initial motor plan *r*(*k*). Notice that in comparison, (7) assumed *u*_*f*_ (*k*) = *r*(*k*), because we had not yet modeled the feedforward system, so there was no learning transfer from the disturbance observer to the feedforward motor plan. In the new model (12), *u*_*f*_ (*k*) is modified through interaction with the disturbance observer so that a correction term *r*_*f*_ (*k*) appears in *u*_*f*_ (*k*), reflecting learning in the feedforward system.

The role of signal *u*_*im*_(*k*), from the point of view of the feedforward system, is to act as a secondary error signal. As shown in our preceding analysis in (8), *u*_*im*_(*k*) asymptotically converges to the negative value of constant perturbations acting on the visual error. If such a perturbation persists over a long time, the disturbance observer must continuously compensate to maintain a properly calibrated visuomotor system. The magnitude of *u*_*im*_ quantifies this ongoing compensation effort. A sustained non-zero *u*_*im*_(*k*) thus informs the feedforward system that it requires a recalibration. In this way, the feedforward system can be viewed as a mechanism for gradually offloading the compensatory burden from the disturbance observer - potentially reducing the biological cost associated with maintaining long-term sensorimotor alignment.

### 3.6 Modeling Assumptions

This section summarizes all modeling assumptions introduced above.

(A1) We assume on the motor side that the transformation from a high level plan to move the arm in a certain direction to the low-level torques applied at the joints is ideal. Mathematically, it means that the parameters of an internal model of the arm in (2a) are matched to the physical parameters of the arm in (2b).

(A2) We assume on the sensory side the subject is equipped with a pre-calibrated visual system so that what is seen in the world is in correspondence with a body-fixed characterization of where is the hand. This assumption means the subject has not put on amplifying or prismatic lenses, etc. The combination of (A1) and (A2) allow the subject to perform baseline trials.

(A3) The brain receives a measurement of the visual error *e*(*k*), and it utilizes an efference copy of the motor command.

(A4) The error sensitivity *K* appearing in the error feedback is a linear function of the error. This assumption is further discussed in Section 4.1.

(A5) The brain contains an internal model of constant perturbation signals to support the visuomotor system. As per the internal model principle, a zero steady-state visual error is therefore achievable for this class of perturbations. Zero steady-state error is no achievable for more complex perturbations such as sinusoids [32].

(A6) The disturbance observer is housed in a part of the brain that supports a recurrent architecture.

(A7) The brain contains a separate feedfoward system for moving the hand when no visual error is present. The feedforward system is calibrated through learning from the disturbance observer.

(A8) All parameters in the model are constants, meaning the model, at present, does not include parameter adaptation laws. This assumption is an outgrowth of prior simulation studies in which we found that the transients generated by the standard gradient law (Widrow-Hoff rule) are incompatible with the transient response of visuomotor rotation experiments [37]. See Section 6.2.

## 4 DO Model

In the previous section we discussed the computational modules that form the building blocks of the DO model. We now face two modeling challenges. First, we must devise a model that correctly updates all computational modules, even when the participant switches between Learn, Ignore, and No Cursor trials. Second, the model must be extended to account for nonlinear saturation behavior during error clamp experiments.

Figures 1-3 show the three modalities that the DO model accounts for. Figure 1 shows that all control modules are active during learning, though they operate on different timescales. Figure 2 shows that only the feedforward system is active during No Cursor trials, whereas the disturbance observer falls into a quiescent state since it receives no error. Figure 3 shows that the feedforward system is likewise active during Ignore trials; however, now the disturbance observer performs silent updates (not expressed in the motor command) because it still receives an error measurement, despite the instructions to ignore the cursor. Our task is to devise pseudo-code that accurately captures the activation or de-activation of control modules for these three scenarios.

To keep track of trials with or without a cursor, we introduce a new index *j* that records the number of trials with a cursor. The software implementation (in Matlab) is structured such that a single value of *j* is associated with each trial number *k* so that *j* may be conceptualized as a function *j*(*k*) of *k*. Since *j* is not updated on each *k* but simply holds its value over No Cursor trials, the plot of *j* = *j*(*k*) as a function of *k* is a staircase function. Notice that because a unique *j* is associated with each *k* (meaning *j*(*k*) is not a multi-valued function), no illegal comparisons between signals indexed with *j* and those indexed with *k* can be made in the pseudo-code implementation below.

Using the two indices *k* and *j, x*(*k*) is the hand angle and *r*(*k*) is the target, both defined for each trial *k*. Instead, *d*(*j*) is the perturbation angle and *y*(*j*) is the cursor angle, both only defined for the *j*-th trial when a cursor is visible. The importance of recording the index separately is that certain signals such as *e*(*j*) will be remembered over No Cursor trials. Without two indices, this memory mechanism is more cumbersome to encode with basic pseudo-code. Finally, all states as well as indices *k* and *j* are initialized to zero (states can be initialized to any value set by the user). The following is the full DO model with switching between experimental modalities.

- If a cursor is displayed on the *k*-th trial, perform the following updates:

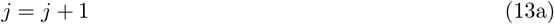

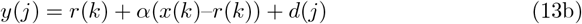

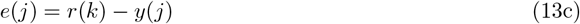

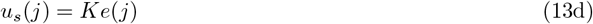

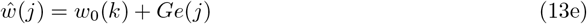

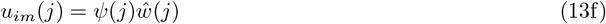

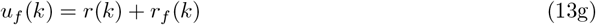

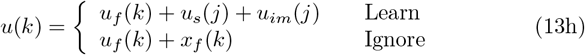

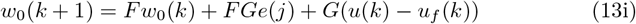

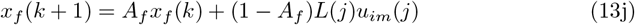

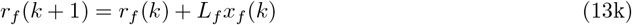

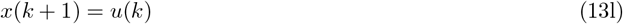
- If no cursor is displayed on the *k*-th trial, perform the following updates:

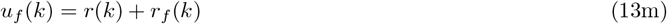

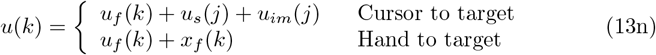

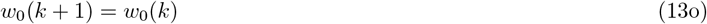

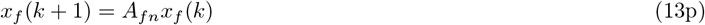

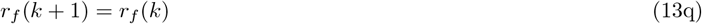

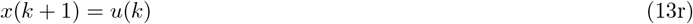

If we select *ψ*(*j*) = *ψ*_o_ and *L*(*j*) = *L*_o_ to be constants, then the DO model is a switched linear state space model. A nonlinear model is obtained by allowing *ψ*(*j*) and *L*(*j*) to vary as a function of the index *j*; see Section 4.1 for this extension. For studies of short-term adaptation, the linear DO model uses five free parameters *K, F, A*_*f*_, *ψ*_o_, and *L*_o_, while the nonlinear DO model uses two additional free parameters *b*_*w*_ and *b*_*f*_. For studies of long-term adaptation, an additional parameter *L*_*f*_ is needed. The role of each parameter and its nominal value are shown in Table 2. The parameters are associated with atomic behaviors of the model. A procedure to select the parameters is given in Section 4.2.

Several features of the DO model warrant examination as they were not already exposed in our discussion of model components. As is indicated, (13a)-(13l) model the computations on trials when the cursor is visible. Eq(13n)-(13r) model the computations performed on trials when no cursor is displayed. The overall motor command used on the next trial is given by (13h). It has two modes, depending on whether the participant is instructed to move the hand to the target or move the cursor to the target on the next trial. Specifically, *u*_*im*_(*j*) is used on trials where the participant is instructed to move the cursor to the target, while *x*_*f*_ (*k*) is used on trials when the participant is instructed to move the hand to the target.

Turning to (13n)-(13r), these equations describe the computations on trials where no cursor is displayed. Since there is no visual error on such trials, the disturbance observer does not update its states; both *w*_0_(*k*) and *ŵ*(*j*) are held constant. This behavior reflects the assumption that participants cannot unlearn a visuomotor perturbation during no cursor trials - unlearning is presumed to require baseline trials with a veridical cursor. However, in re-analyzing data from two previously published experiments, we found that incorporating a small forgetting factor (*F*_*n*_ = 0.95) for *w*_0_(*k*) for no cursor trials improved the DO model’s fit to participant behavior. This model extension will be discussed in Section 6. While the disturbance observer nominally remains quiescent during no cursor trials, the feedforward system continues to update in (13p)-(13q). Specifically, *r*_*f*_ (*k*) is held constant, whereas *x*_*f*_ (*k*) decays at a rate determined by parameter *A*_*fn*_. As such, *A*_*fn*_ is the sole parameter that determines the decay rate in No Cursor trials when the subject moves the hand to the target.

The DO model characterizes mathematically how the brain manages three experimental modalities in visuomotor experiments: Learn, Ignore, and No Cursor trials, where No Cursor trials are associated with moving the hand to the target. We see that the DO model has a fourth modality in (13n) to move the cursor to the target on the next trial after a No Cursor trial. This modality has been included to capture behaviors in experiments involving rapid switching between Learn and No Cursor trials in [18, 38]. Effectively, the error is remembered from the previous trial it was visible so that the subject retains the ability to perform a Learn trial despite there being no cursor visible on the immediately preceding trial.

The DO model is structured so that the disturbance observer’s estimate of the perturbation does not depend on the specific mode of the motor command *u*(*k*); whether the participant is instructed to move the hand or the cursor to the target, the disturbance observer still converges to an accurate estimate of the perturbation. This behavior can be understood from the calculation in (8) where we saw that *u*(*k*) is canceled in the update of *ŵ*(*k*). The same behavior remains true for updates of *ŵ*(*j*) using the index *j* to record cursor trials only. Furthermore, if we repeat the calculation in (8) for a constant perturbation *d*(*j*) = *d*, but we allow for non-zero values of *r*_*f*_ (*k*), then we get

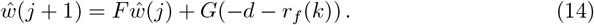

Note again that the comparison between indices *j* and *k* is consistent if we interpret *j* as a function of *k*. This calculation shows that the amount of the perturbation that must be accounted for by the disturbance observer is adjusted by the feedforward correction *r*_*f*_ (*k*). If the adjustment in *r*_*f*_ (*k*) converges to −*d*, then asymptotically *ŵ*(*j*) evolves according to

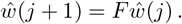

Since this is a stable linear system, it means over the long term, *ŵ*(*j*) tends to zero. The implication is that over the long term, the feedforward system would be capable to fully offload the work of the disturbance observer.

For the feedforward system to operate properly, the *learning transfer rate L*_*f*_ should be sufficiently small. Formally speaking, this constraint gives rise to a two timescale control architecture [39]. The overall operation of the two timescale system goes as follows. When an environmental perturbation occurs, it is counteracted by the first responder available: an error feedback *u*_*s*_(*j*). If the perturbation persists, then a somewhat slower but still short-term process is activated in *ŵ*(*j*) to estimate and eliminate constant perturbations. As the disturbance observer output *u*_*im*_(*j*) accumulates information about the perturbation, this drives the activation of the state *x*_*f*_ (*k*). The transfer of information from *u*_*im*_(*j*) to *x*_*f*_ (*k*) is subject to an error sensitivity *L*_o_ (or a strong saturation (15b) defined below), which constrains the extent of learning at this stage. A sustained positive or negative *x*_*f*_ (*k*) drives the adaptation of *r*_*f*_ (*k*). The role of *r*_*f*_ (*k*) is to incorporate the history of a persistent perturbation into the feedforward motor output, modifying it to the form *u*_*f*_ (*k*) = *r*(*k*) + *r*_*f*_ (*k*). The latter expression represents an updated motor command that reflects prolonged exposure to the visuomotor perturbation.

A most remarkable property of the DO model is that *r*_*f*_ (*k*) will asymptotically estimate − *d* in the long term. As such, *r*_*f*_ (*k*) represents the system’s long-term memory of the perturbation that becomes “hardwired” into the feedforward motor command. A full analysis of this remarkable behavior will be left for a future study that also provides supporting experiments of long-term adaptation.

### 4.1 Error Sensitivity Functions

The DO model includes three parameters *K, ψ*(*j*), and *L*(*j*), each of which may be interpreted as error sensitivity functions. The first, *K* corresponds to single trial or trial-to-trial learning [27, 28]; reflecting the system’s immediate sensitivity to the observed visual error *e*(*j*). Empirical studies have shown that this sensitivity decreases nonlinearly for larger errors, deviating from a strictly linear regime [27–29]. In the present model, this nonlinearity has not been implemented, so the error sensitivity function is a linear function of the error. The second error sensitivity function *ψ*(*j*) accounts for *incomplete learning*, a phenomenon consistently observed in visuomotor adaptation tasks [21, 40–42]. The third function *L*(*j*) represents the sensitivity to the secondary error *u*_*im*_(*j*), and it plays a key role in shaping the system’s response under error clamp conditions [22, 25]. By modulating the gain on this pathway, *L*(*j*) enables the DO model to replicate the plateauing of adaptation observed when the visual error is held constant.

We have selected the nonlinear error sensitivity functions to be:

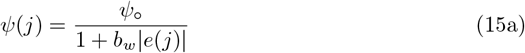

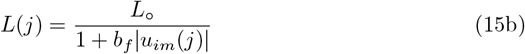

These two functions are the only parts of the model that are adhoc. Function *L*(*j*) was constructed to best fit the saturation observed in the graded error clamp of Experiments 1-4. Function *ψ*(*j*) simply mimicked the same structure as *L*(*j*). It’s presence is for closed-loop stability during non-zero error clamp trials. Further experimental research is needed to verify the accuracy of these formulas, whose role is to capture safety mechanisms in the brain.

### 4.2 Selection of Parameters

We recommend an ordered procedure to select the parameters of the DO model consisting of three steps. Table 2 summarizes the parameters, their nominal values, and their functional roles. Additional guidance and empirical justification are provided in the Results section.

1. **Parameters for the disturbance observer**. The first step is to select parameters that govern the classical behaviors of visuomotor adaptation: savings with counter perturbation, spontaneous recovery, and anterograde interference. These behaviors are particularly well-documented for the saccadic system [43] and in force field experiments [44, 45]. In the DO model these behaviors are captured by a linear disturbance observer with three free parameters: *K, F*, and *ψ*_o_.
  - Parameter *K* sets the gain for single trial learning. Across the experiments we considered, *K* = 0.25 is a reasonable nominal value, including for the saccadic system and visuomotor adaptation. This parameter is responsible for the inflection that was observed for the saccadic system and identified with arrows in Figure 3A of [43]. By making *K* larger, this inflection is made more pronounced, amplifying sensitivity to early errors.
  - Parameter *F* sets the learning rate of the disturbance observer and must lie in the interval (0, 1). It roughly corresponds to the learning rate of the slow process in a two-rate model (a coordinate transformation is needed to make the precise comparison). Our observation is that for visuomotor experiments with relatively fast learning, one can set *F* = 0.7. For the saccadic system as well as for force field experiments, the response is somewhat slower, so a value of *F* = 0.9 is more suitable. To accentuate the two rate classical behaviors of visuomotor adaptation, one should select a larger value of *F* = 0.9 to better separate the timescales in the two-rate model. Note that *G* = 1 − *F*, so it is not a free parameter.
  - Parameter *ψ*_o_ sets the fraction of learning at asymptote. When a participant fully compensates for the perturbation, then *ψ*_o_ = 1. However, many studies report incomplete learning [21, 40–42]. In this case a value of *ψ*_o_ = 0.95 is typical.
  Once the parameters of a standard learning paradigm have been identified for a given experiment, these parameters should not be modified, as they fully characterize standard learning.
2. **Parameters for the feedforward system**. The feedforward system is active in trials with no cursor, zero error clamp trials, and trials with instructions to move the hand to the target (while ignoring the misaligned cursor). The linear feedforward system utilizes two parameters: *A*_*f*_ and *L*_o_.
  - Parameter *A*_*f*_ determines the learning rate of the feedforward system. Based on analysis across multiple datasets, a nominal value is *A*_*f*_ = 0.9 provides a good approximation. However, there can be some variability in this parameter, which we report on below.
  - The parameter *L*_o_ is the error sensitivity to the secondary error signal *u*_*im*_(·). This parameter affects the steady state response of the feedforward system and is particularly relevant under error clamp conditions. It is best to calibrate *L*_o_ using small error clamp values within the linear range of system response; see [25].
3. **Nonlinear model parameters**. For experiments involving non-zero error clamp trials in the nonlinear zone, it is necessary to introduce saturation in two areas of the DO model.
  - Parameter *b*_*f*_ controls the saturation behavior of the feedforward system under error clamp conditions. Specifically, it governs the nonlinear attenuation of steady state responses to sustained error signals. Increasing *b*_*f*_ amplifies the saturation effect, thereby reducing the asymptotic output of the feedforward system. Because this parameter significantly shapes the DO model’s behavior in the nonlinear regime, it should be used judiciously and only to capture empirical patterns specific to high-error clamp conditions. Notably, there is variability in reported steady state responses across studies: [25] documents higher asymptotic values than those observed in the similar experiment of [23]. As such, tuning *b*_*f*_ requires careful calibration against the particular experimental dataset being modeled.
  - Parameter *b*_*w*_ constrains the growth of signals in the disturbance observer during non-zero error clamp trials. Without this constraint, the linear disturbance observer can produce unbounded growth of signals. Introducing nonlinear saturation via a small value of *b*_*w*_ = 0.001 effectively prevents divergence while leaving standard adaptation behavior unaffected. This ensures the DO model remains stable during prolonged clamp conditions without distorting its performance under typical learning paradigms.

### 4.3 Adaptation Fields

It is well known that the visuomotor system is able to perform dual adaptation, namely adaptation to two or more perturbations in the visual field [46–51]. The phenomenon is particularly evident when perturbations are associated with separate workspaces or targets [47–49]. The DO model presented here regards a single target and a single direction of the perturbation (CW or CCW). If one is interested to capture dual adaptation using the DO model, it is necessary to create separate computational modules for each adaptation field. Particularly, each adaptation field requires a dedicated disturbance observer to learn independent perturbations.

## 5 Results

We conducted a new experiment and also drew on datasets from both within and outside our lab to evaluate how well the DO model captures a range of phenomena in visuomotor adaptation. The first three sections - spontaneous recovery, savings, and the error clamp – address well-established behaviors and are therefore described briefly. Sections 5.4–5.7 focus on more complex behaviors that test the DO model’s explanatory power, either through novel predictions or by accounting for effects not captured by other models. The *reach deviation* depicted in the figures is defined as *x*(*k*) − *r*(*k*). For consistency, simulation plots show (primarily) positive hand angles to facilitate comparison.

### 5.1 Spontaneous Recovery

Spontaneous recovery is a rebound phenomenon that can be invoked in either washout, zero error clamp, or no cursor trials following a sufficiently long phase of Learn trials with a short counter-perturbation phase. Fig 7(a) depicts the classical mechanism of spontaneous recovery induced by a short counter-perturbation followed by a washout phase. This behavior is elicited in the DO model using only the disturbance observer. The behavior arises from a separation of timescales between slow and fast processes, which can be accentuated by slowing the disturbance observer learning rate to *F* = 0.9. Fig 7(a) shows the trace for *ŵ*(*j*), demonstrating that spontaneous recovery arises because the short counter-perturbation phase does not allow the slow component of learning *ŵ*(*j*) to fully decay. Fig 7(b) illustrates evoked recovery [45], a more sophisticated form in which recovery emerges in No Cursor trials, driven by the feedforward system. The trace of *r*_*f*_ (*k*) + *x*_*f*_ (*k*) demonstrates that the output of the feedforward system is responsible for this form of spontaneous recovery. This signal is hidden during Learn trials which are driven by the disturbance observer, but emerges at the transition from Learn to No Cursor trials. This experiment highlights that spontaneous recovery can arise either from behavior of the disturbance observer or the feedforward system, depending on the context.

**Fig 7.**
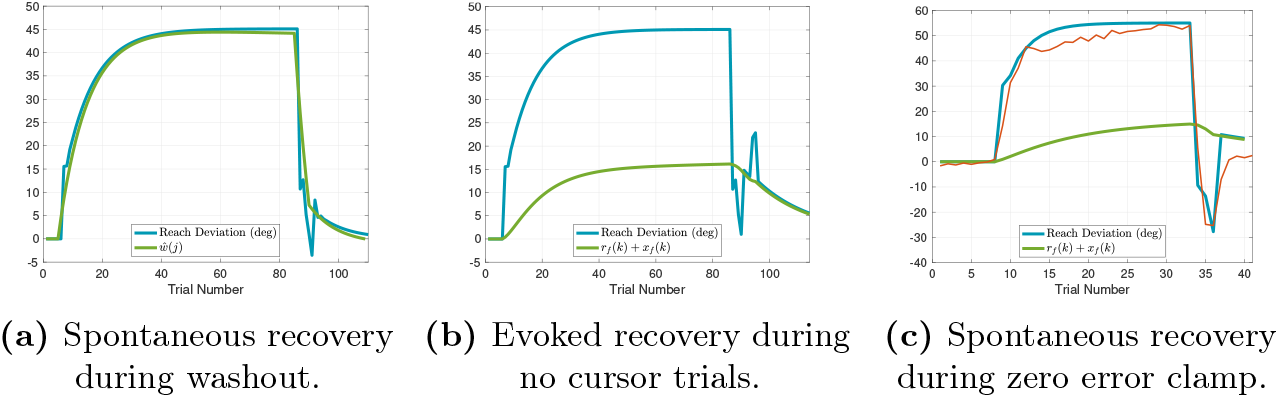
Spontaneous Recovery. (a) Spontaneous recovery in an experiment of 80 Learn trials with a − 45° perturbation, 5 unlearning trials with a +45° perturbation, and 20 washout trials with veridical feedback. The green trace of *ŵ*(*j*) shows that the disturbance observer is responsible for spontaneous recovery in this experiment. (b) Evoked recovery in an experiment of 80 Learn trials with a −45° perturbation, 4 unlearning trials with +45° perturbation, 3 no cursor trials, 2 Learn trials with a −45° perturbation, and 20 no cursor trials. The green trace of *r*_*f*_ (*k*) + *x*_*f*_ (*k*) shows that the feedforward system is responsible for spontaneous recovery in this experiment. (c) Spontaneous recovery in an experiment from [52] of 25 Learn trials with a +60° perturbation, 3 unlearning trials with a − 60° perturbation, and 5 zero error clamp trials. The red trace is the experimental data and blue traces are model predictions. The feedforward system is again responsible for spontaneous recovery in this experiment. Recall that hand angles are shown with positive values (primarily) for ease of comparison.

Fig 7(c) demonstrates another variant - spontaneous recovery elicited in zero error clamp trials - with simulation results compared to the data from [52]. To model zero error clamp trials, we follow the proposal of [53] that a brain process actively disengages at the transition from Learn trials to zero error clamp trials. Indeed, the error may be small at the end of a long sequence of Learn trials, yet when the zero error clamp is enforced, [53] shows that the hand angle decays. Additional evidence from [54] shows similar forgetting rates between no cursor trials and zero error clamp trials. Accordingly, we model zero error clamp trials as driven by the feedforward system, resulting in computations that are equivalent to no cursor trials.

### 5.2 Savings

Savings in the visuomotor system is a mechanism to perform recurring visuomotor tasks with improved performance and lower biological cost. It should be no surprise that savings is a multifaceted phenomenon, reflecting a multiplicity of strategies to achieve this desirable behavior, an idea supported by a range of sometimes conflicting experimental findings [55–58]. The DO model can reproduce at least four distinct forms of savings: (i) classical savings using counter-perturbations [44]; (ii) savings due to re-aiming [57]; (iii) savings due to increased error sensitivity [59]; and (iv) savings via learning transfer, which is addressed in Section 5.7. Figure 8 illustrates simulations of the first three mechanisms, with comparisons to experimental data from [56].

**Fig 8.**
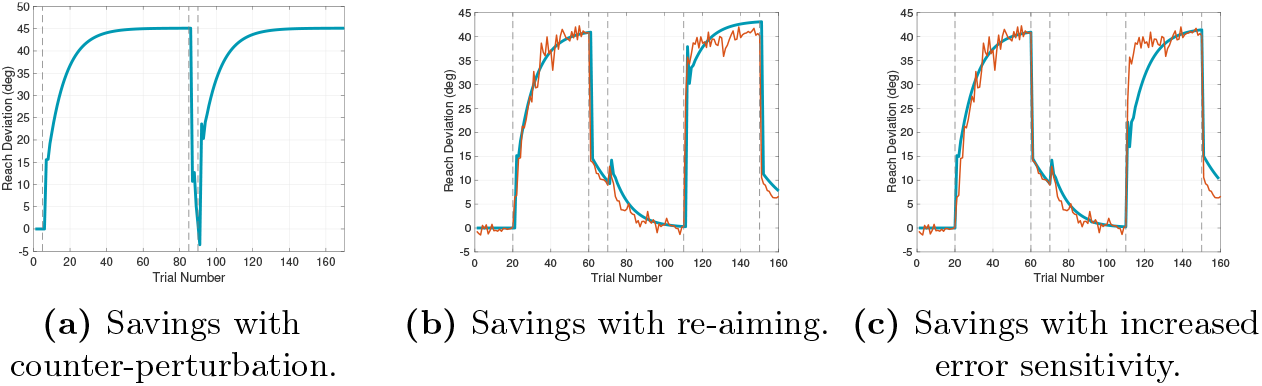
Savings. (a) Savings in an experiment of 80 Learn trials with a − 45° perturbation, 5 unlearning trials with a +45° perturbation, and 80 relearning trials. (b) Savings with re-aiming in Experiment 1 of [56] with 40 Learn cycles (4 targets per cycle) with a − 45° perturbation, 10 no cursor cycles, 40 washout cycles, and 10 no cursor cycles. (c) Savings with increased error sensitivity in Experiment 1 of [56]. Red traces correspond to experimental data and blue traces correspond to model predictions.

Fig 8(a) shows the classical notion of savings in a paradigm involving a brief counter-perturbation phase. The faster relearning in the second exposure to the perturbation arises because the disturbance observer retains information from the first learning phase. Due to its slower dynamics relative to the fast component *e*(*k*), it partially retains this knowledge over the brief counter-perturbation. This savings mechanism is in accordance with the predictions of a two-rate model [44]. Fig 8(b) shows the effect of re-aiming on savings in Experiment 1 from [56]. Re-aiming is implemented in the second learning phase by setting *r*_*aim*_(*k*) = − *γd* with *γ* = 0.5, where *γ* ∈ (0, 1) specifies the proportion of the perturbation compensated through re-aiming. Fig 8(c) shows the effect of increased error sensitivity on savings. Increased error sensitivity is implemented in the DO model by increasing the model’s error sensitivity parameter to *K* = 0.5 during the washout and second learning phases. The notion that the visuomotor system can adjust *K* is plausible and consistent with a *gain control mechanism* extensively investigated for the saccadic and smooth pursuit eye movement systems [60, 61].

### 5.3 Error Clamp

The error clamp paradigm offers a platform to explore nonlinear saturation in visuomotor adaptation. It has been observed over a number of studies that the amount of saturation at asymptote can be inconsistent [22, 23, 25]. The authors of [23] speculate this variability may stem from differences in experimental design - such as the use of endpoint versus continuous cursor feedback, or due to blanking the target after 250ms. While the DO model does not directly account for such experimental conditions, two parameters associated with the nonlinear feedforward system *L*_o_, and *b*_*f*_ can be tuned to reproduce the range of behaviors observed under the error clamp. We use the data from [23] to illustrate the idea. Fig 9(a) compares the hand angle across three conditions: standard movement contingent learning, learning with instructions to move the hand to the target, and the error clamp with instructions to move the hand to the target; compare to Fig 1C of [22]. Fig 9(b)-(c) show simulations in the linear and saturated regimes of the error clamp; matching the behavioral profiles shown in Figs 1c-d of [25]. Fig 9(d) replicates the more suppressed response observed in [23]; this requires increasing the value of *b*_*f*_, suggesting that this parameter is key to matching response magnitudes across different error clamp studies.

**Fig 9.**
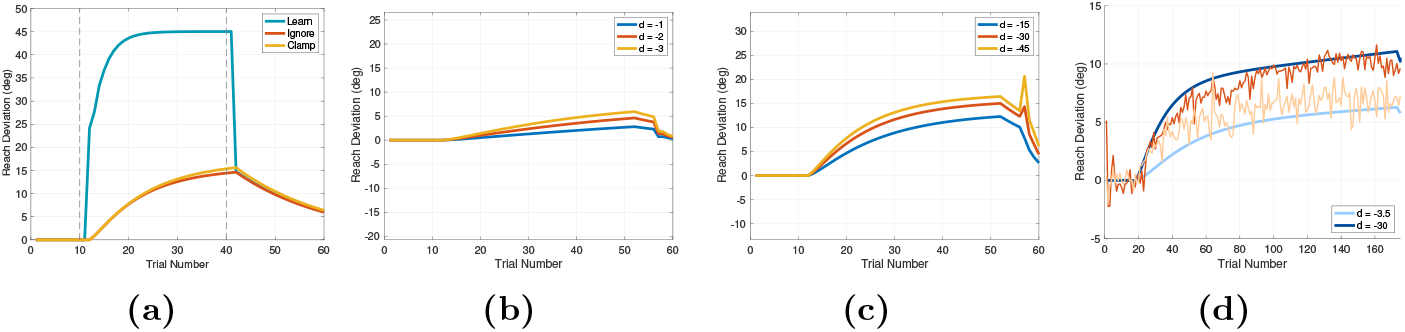
Error Clamp. (a) Experiment 1 in [22] comparing Learn trials with a − 45° perturbation and instructions to move the cursor to the target; Ignore trials with instructions to move the hand to the target; and error clamp trials with a clamp size of *d* = − 45° with instructions to move the hand to the target. (b)-(c) Error clamp in the proportional (b) and saturated zones (c); see Figs 1c-d of [25]. (d) Experiment 1 of [23] using a cursor.

The DO model also accounts for error clamp behavior in cerebellar patients. In Experiment 3 of [22], a group of 10 participants with varying degrees of cerebellar ataxia performed an experiment with a 45° error clamp and instructions to move the hand to the target. Their responses (see Fig 3A of [22]) were clearly attenuated compared to controls. This outcome can be explained using the DO model by the fact that the feedforward system, which drives the computations when moving the hand to the target, requires as input *u*_*im*_ from the disturbance observer. We hypothesize the disturbance observer resides in the cerebellum (see the last section of the paper), so this output signal will either be absent or reduced in cerebellar patients, leading to diminished feedforward responses in the error clamp condition.

### 5.4 Graded Error Clamp

A central challenge in modeling visuomotor adaptation is to understand if the saturation behavior of the error clamp is an isolated phenomenon, a *singularity*, or if it is a ubiquitous phenomenon in all visuomotor tasks. To investigate this question, we designed a new experimental paradigm called the **graded error clamp**. This technique incrementally introduces error clamp conditions by adjusting a single parameter *α* which scales the contribution of the hand angle to the displayed cursor position. While similar scaling approaches have appeared in prior studies under the term “magnification of errors” [20, 21] (compare Eq 2 in [21] with our (13c)), to our knowledge this method has not previously been used to *gradually implement* the error clamp.

In our study, each participant performed the Ignore-N and Learn-N experiments in two sessions. As expected, we found no significant main effect of session or interaction between session and the other factors (*F* (1, 29) = 0.07, *p* = 0.80). Therefore, responses were averaged over the two sessions. During Ignore trials in the Ignore-N experiment, participants experienced a rotation of −15° for *α* ∈ {1.0, 0.6} and +15° for *α* ∈ {0.8, 0.4}. The same rotation values were used in the Learn-N experiment during Learn trials. The design is summarized in Fig 6.

Simulations corresponding to these two experiments are shown in Fig 10 superimposed with the experimental data. Data were separated by perturbation direction (CW and CCW) due to observed asymmetries in responses, particularly in the Learn-N experiment. In the Ignore-N experiment with instructions to move the hand to the target during Ignore trials, the DO model predicts that only the feedforward system makes a direct contribution to the hand movement. The response is suppressed relative to Learn-N experiment with the corresponding *α* value, as one would expect. According to the model, this suppressed response is due to the graded error clamp condition, which causes the nonlinear error sensitivity *L*(*j*) to suppress the response of the feedforward system.

**Fig 10.**
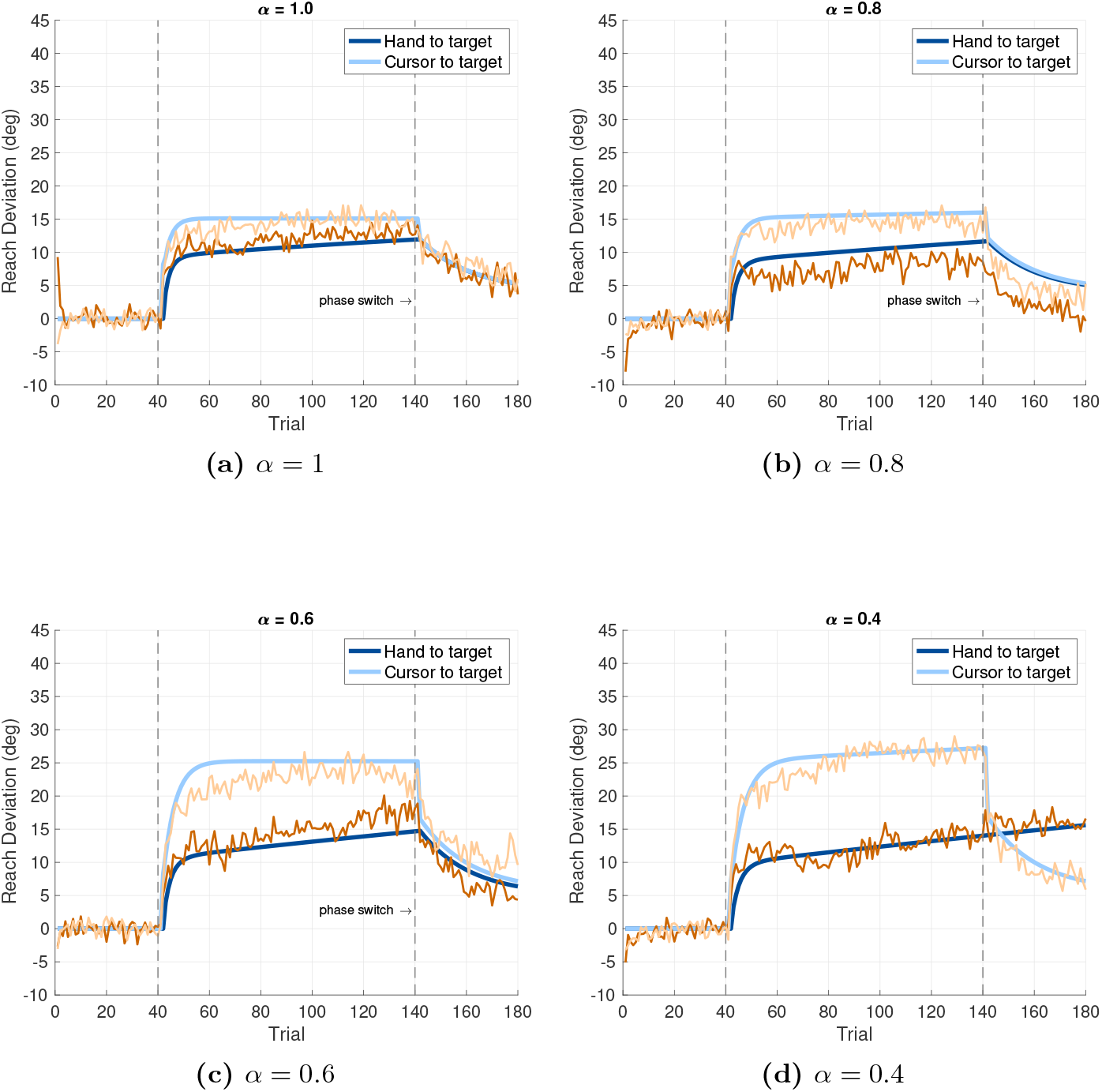
Graded Error Clamp. Simulation compared to experimental data in Experiments 1 and 2. Experiment 1 consists of 40 baseline trials, 100 Ignore trials, and 40 no cursor trials. For *α* = 0.4, the number of Ignore trials is 160. Experiment 2 consists of 40 baseline trials, 100 Learn trials, and 40 no cursor trials. The perturbation is −15° for *α* ∈ {0.6, 1.0} and +15° for *α* ∈ {0.4, 0.8}. A phase switch is predicted to occur at the transition from Learn to No Cursor trials. Orange traces correspond to experimental data and blue traces correspond to model predictions.

In the Learn-N experiment, the DO model predicts that the hand movement is driven by the disturbance observer. The feedforward system state *x*_*f*_ (*k*) does not make a direct contribution during these trials, but becomes active during subsequent No Cursor trials. To achieve zero steady state error in the Learn trials, the theoretically required steady state value of the hand angle is

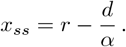

With the values *r* = 90°, *d* = −15, and *α* ∈ {0.4, 0.6, 0.8, 1.0}, we expect (*x*_*ss*_ −*r*) ∈ {37.5, 25, 18.75, 15}. These steady state values of the hand angle were observed experimentally in Experiment 2 for negative (CW) perturbations *d* = − 15 corresponding to *α* ∈ {1.0, 0.6}. However, for *α* ∈ {0.4, 0.8} and positive (CCW) perturbations of *d* = +15, we found participants did not fully compensate for the perturbation. Recalling that the parameter *ψ*_o_ is the only parameter in the DO model to capture the proportion of learning at asymptote, it was necessary in the simulations to reduce the parameter *ψ*_o_ in the DO model to a value *ψ*_o_ = 0.8 (from its nominal value of *ψ*_o_ = 1) in order to capture incomplete learning observed experimentally. This difference may reflect a bias in visuomotor adaptation where arm movements away from the saggital plane are suppressed due to the risk of generating inappropriate torques on the shoulder and elbow joints. The parameter *ψ*_o_ may be viewed as a proxy for this safety mechanism.

To better understand the results of the first two experiments, we conducted Experiments 3 (Ignore-W) and 4 (Learn-W), in which each participant only experienced one perturbation direction: either CW or CCW. This eliminated the possibility of carry-over between phases of the experiments. We found that the pattern of incomplete learning persisted for participants who experienced CCW perturbations (a value of *ψ*_o_ = 0.8 again captures this incomplete learning in the DO model). Figure 11 shows the experimental results averaged over CW and CCW perturbations. We used a value of *ψ*_o_ = 0.9 in the DO model, corresponding to averaging responses with *ψ*_o_ = 0.8 (CCW) and *ψ*_o_ = 1 (CW).

**Fig 11.**
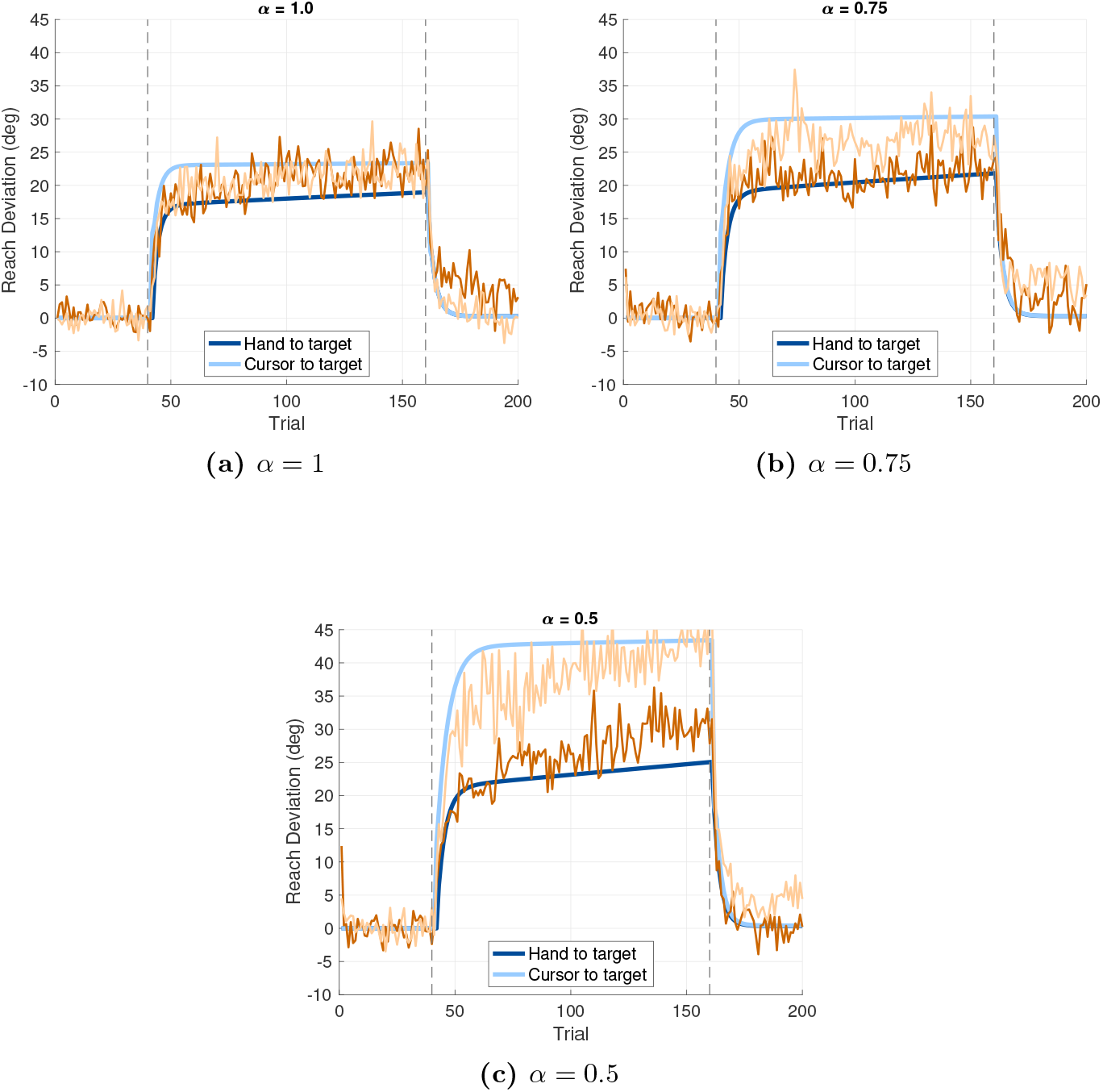
Graded Error Clamp. Simulation compared to experimental data in Experiments 3 and 4. A phase switch is predicted to occur at the transition from Ignore to washout trials. Orange traces correspond to experimental data and blue traces correspond to model predictions.

### 5.5 Phase Switch Behavior

A **phase switch** is defined to be a switch in the *actively expressed* computational module or stream in the brain during visuomotor adaptation. The DO model includes two distinct computational modules that perform updates concurrently: a disturbance observer and a feedforward system. However, only one of these two processes is actively expressed in the motor command on any trial (as seen in (13h)). The DO model proposes that a phase switch occurs at the following experimental transitions:

- Learn trials to Ignore trials, and vice versa.
- Learn trials to no cursor trials without explicit instructions, where the absence of a cursor leads participants to default to moving their hand to the target.
- Learn trials to zero error clamp trials, as supported by findings in [53].

The concept is illustrated using the data from [58]. Fig 12(a) shows their control group experiment consisting of 40 Learn trials, 10 no cursor trials, 20 washout trials, and 40 relearning trials. The DO model predicts two phase switches: one at the transition from Learn to no cursor trials, and the second from no cursor to washout trials. Fig 12(b) shows the clamp group experiment consisting of 40 error clamp trials, 10 no cursor trials, 20 washout trials, and 40 Learn trials. Here the DO model predicts only one phase switch at the transition from no cursor to washout phases. We note that the prediction of phase switches in experiments such as [58] is not based on what we see in the data, but based on a modeling hypothesis that the brain makes a switch in computational streams for reasons such as a change in instructions. The hypothesized existence of phase switches must be further tested, potentially with experiments involving brain imaging or noninvasive brain stimulation.

**Fig 12.**
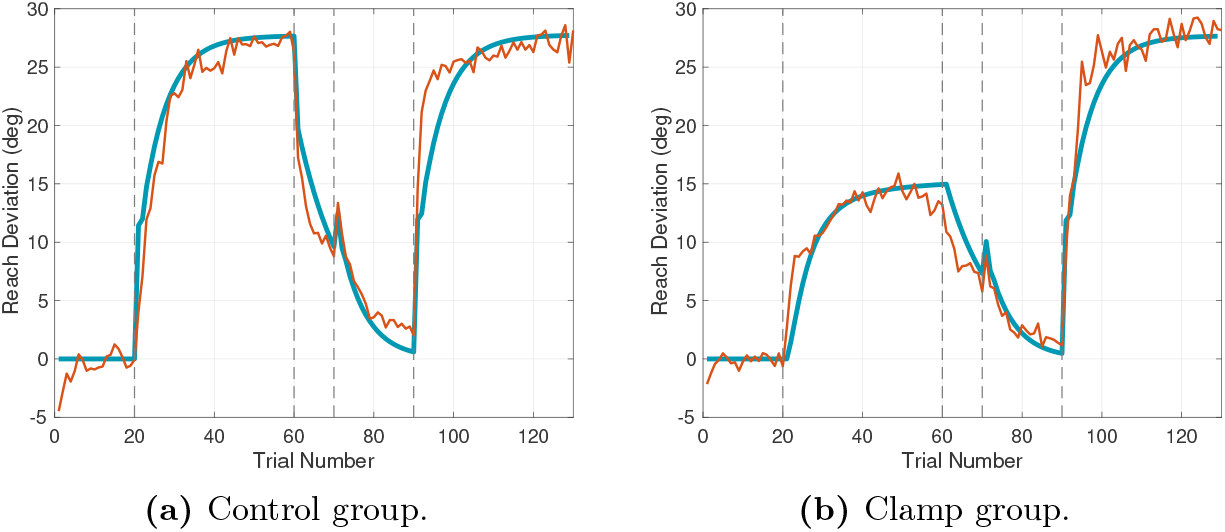
Phase Switch. Demonstration of phase switches using Experiment 1 of [58]. Orange traces correspond to experimental data and blue traces correspond to model predictions.

Experiments 1-4 also demonstrate behavior attributable to a phase switch. In the Learn-N Experiment a phase switch is predicted to occur at the transition from Learn trials to no cursor trials, resulting in the characteristic drop in the hand angle, particularly visible for *α* = 0.4. In the Ignore-N Experiment no phase switch is expected between Ignore and no cursor trials, leading to a smoother transition. In the Ignore-W Experiment, a phase switch is predicted between Ignore and washout trials, whereas the Learn-W Experiment is predicted to have no phase switch between Learn and washout trials. In Experiments Ignore-W and Learn-W the discrepancy between a phase switch and no phase switch is more difficult to discern, likely because we used washout trials with veridical cursor feedback rather than no cursor trials, as in the Ignore-N and Learn-N Experiments.

One of the most compelling demonstrations of a phase switch is an experiment reported in [62]. In their Experiment 3, participants performed a sequence of Learn trials with a perturbation size *d* ∈ {15, 30, 45, 60, 90}, with one perturbation size for each group of participants. Following the Learn trials, participants performed a series of no cursor trials with instructions to move the hand directly to the target. It was shown in Fig 7A of [62] that aftereffect sizes in the no cursor phase did not significantly differ between groups who experienced different rotation sizes. The DO model accounts for this counterintuitive result by proposing the involvement of a hidden process during the Learn phase that becomes observable only after a phase switch at the onset of no cursor trials. Fig 13(a) presents simulations of this experiment, replicating the pattern reported in Fig 7A in [62]. The interpretation based on the DO model is that the hidden process is the feedforward system - it performs silent updates during Learn trials that are driven by the disturbance observer. These updates are not expressed behaviorally until the phase switch occurs, at which point the computations of the feedforward system are revealed.

**Fig 13.**
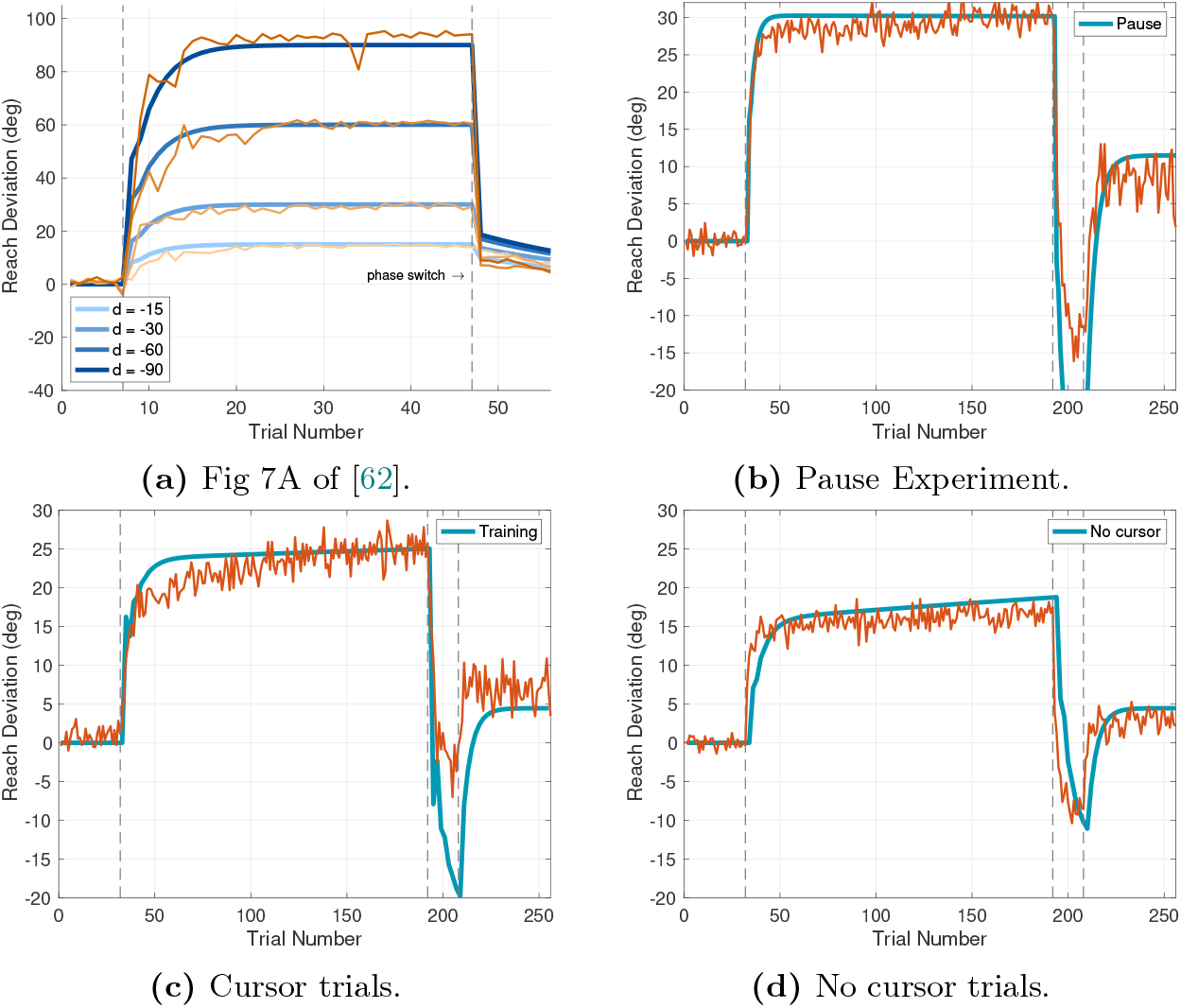
Phase Switch. (a) Experiment 1 of [62]. (b) Pause Experiment of [18] with a brief pause after every Learn trial. (c) Cursor - No Cursor Experiment of [18], alternating Learn and no cursor trials. On odd-numbered Learn trials, participants were instructed to move the cursor to the target with the cursor visible. (d) On even-numbered no cursor trials, participants were instructed to move the hand to the target with no cursor. The perturbation size is 30°. Orange traces correspond to experimental data and blue traces correspond to model predictions.

Another striking demonstration of phase switch behavior comes from the experiment reported in [18] from our lab. This experiment belongs to a class of paradigms in which participants alternate between two types of trials: (1) moving a rotated cursor to the target with visual feedback of the cursor, and (2) moving the unseen hand to the target with no cursor. The trial types alternate systematically — 30° rotated cursor trials followed by no cursor trials. This design is intended to measure implicit learning expressed during no cursor trials, arising from exposure to the rotated cursor during the preceding visual feedback trials.

We hypothesize that this experiment induces a form of phase switching that continuously engages two distinct computational streams in the brain, each receiving input that conflicts with the subsequent trial’s task demands. Given this conflict, it is not suprizing that some degree of interference arises - particularly when compared to standard 30° Learn phases that do not include interleaved no cursor trials (Fig 1A in [18]). Fig 13(b)-(d) show simulation results corresponding to the experiment in [18]. In Fig 13(b) participants perform a sequence of Learn trials, each followed by a brief pause. According to the DO model, only the disturbance observer is engaged, so there are no phase switches during the Learn and Unlearn phases of the experiment. We see that participants are able to compensate for the full 30° perturbation. Fig 13(c)-(d) provide evidence of interference between two processes. We see in Fig 13(c) that on cursor trials, the disturbance observer is not fully able to learn the 30° perturbation. Meanwhile, the feedforward system (whose output is depicted in Fig 13(d)) achieves a steady state around 15°, resembling behavior observed in error clamp conditions. This occurs because the feedforward system experiences an *effective error clamp* because the disturbance observer fails to vanquish the visual error. The effective clamped error in cursor trials is indistinguishable in the DO model from an experimentally imposed clamp condition, even though the former arises from a natural phenomenon of incomplete learning. The experiment further supports the view that the error clamp condition is a ubiquitous phenomenon that the feedforward system must continuously contend with under reasonably natural conditions.

We note that the DO model does not well-match the experimental data in the counter-perturbation phase of both the pause experiment and the cursor-no cursor experiments, as seen in Fig 13(b)-(c). Finally, Fig 14 shows simulation results for the experiment in [38] which has the same structure as the experiment in [18] but uses values of *d* ∈ {+15, +30, +45, +60} (with *α* = 1).

**Fig 14.**
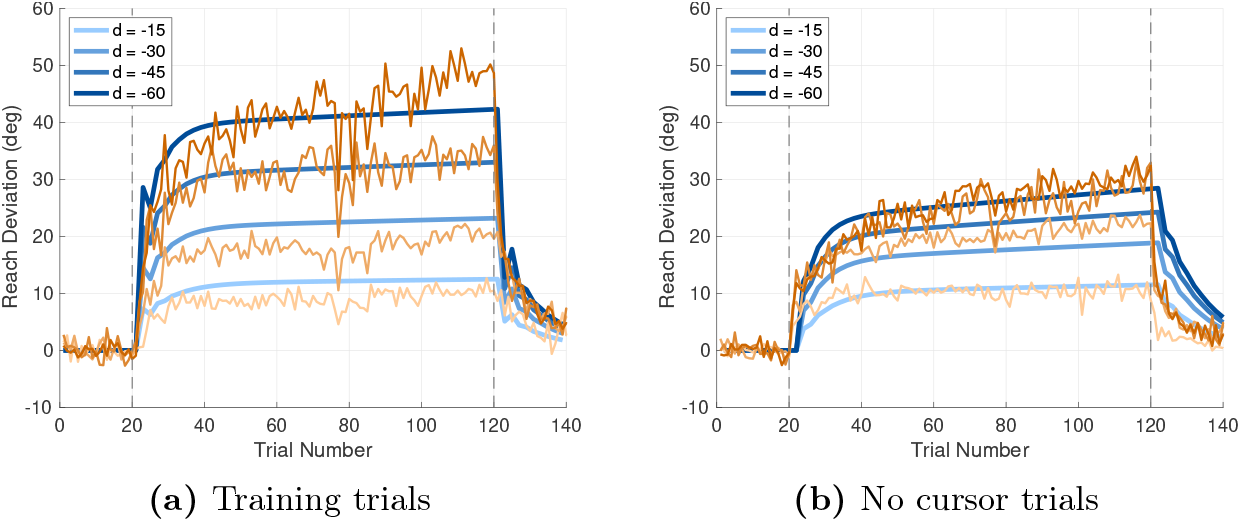
Simulation of the experiment in [38] with rapid switching between cursor and no cursor trials. As in the study of [18], on (a) on odd-numbered training trials, participants are instructed to move the cursor to the target while the cursor is visible during and at the end of the reach; while (b) on even-numbered trials, participants are instructed to move the hand to the target while the cursor is not visible. In contrast with [18], the perturbation size is varied and also the length of the experiment is extended. The perturbation size for the training trials is shown on both figures. Orange traces correspond to experimental data and blue traces correspond to model predictions.

### 5.6 Explicit and Implicit Computations

The DO model may be utilized to analyze explicit and implicit contributions to visuomotor adaptation, a subject of significant interest in visuomotor learning [63]. Since the literature includes several interpretations of the terms “implicit” and “explicit”, here we focus on an interpretation that the explicit part of adaptation regards that part of the motor command over which the participant has volitional control. The DO model suggests that the division between implicit and explicit components is somewhat more subtle than what is currently conceived. If we split the computations into two processes (the disturbance observer and the feedforward system), then the process whose computations are silent over a sequence of trials may appear to be implicit, even if those computations are revealed at a later phase in the experiment and can be altered by the participant.

In order to untangle this situation, we separately analyze Learn trials versus Ignore trials. We assume the number of trials is sufficiently small so that we can take *r*_*f*_ (*k*) ≡ 0. First, consider Learn trials when the participant is instructed to move the cursor to the target. A division of the motor command between explicit and implicit computations is

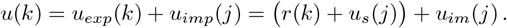

The feedforward command *r*(*k*) is explicit because the participant has efficacy to choose to reach to a specific target. The error feedback *u*_*s*_(*j*) = *Ke*(*j*) is also explicit based on an interpretation that the participant deliberately aims in the opposite direction of the perturbation by an amount that is a fraction of the observed error in the last reach. The computations of the disturbance observer are regarded as implicit, such that the participant has no authority to modify these computations - they progress automatically in a “machine-like” way.

This division follows the traditional view of implicit and explicit computations using a two-rate model. The implications will therefore be familiar. First, the explicit component makes a larger contribution at the beginning of learning when errors are large. The implicit component makes a relatively larger contribution near the end of learning when errors are small. Second, the explicit component makes a larger contribution when the perturbation is larger. Third, the explicit component has no aftereffects associated with it. Any aftereffects would be entirely attributable to *u*_*im*_(*j*) (we are speaking only about aftereffects following Learn trials, with the cursor always visible). Finally, if increased reaction time is associated with the explicit component, and increased aftereffect is associated with the implicit component, then certain experiments will result in a correlation between increased reaction time and decreased aftereffect.

Next, consider Ignore trials when the participant is instructed to move the unseen hand to the target. We propose that the motor command be split into explicit and implicit components as follows:

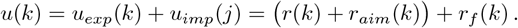

The component *r*(*k*) + *r*_*aim*_(*k*) is explicit because the participant still has the efficacy to choose to reach for a specific target as well as to use an aiming strategy. The component *r*_*f*_ (*k*) generated by the feedforward system is implicit in the sense that this component cannot be suppressed by the participant, even if it results in movements that do not seem useful for the task (moving the hand to the target). Notice that when probing trials are used in visuomotor reach experiments, based on the available data, we believe participants are reporting the full motor command *r*(*k*) + *r*_*aim*_(*k*) + *r*_*f*_ (*k*).

Participants may be aware of the implicit component *r*_*f*_ (*k*) in their reporting, but they do not have the volition to alter or remove it.

Using these ideas, we may consider an experiment in which participants are instructed to deploy an aiming strategy based on information about the perturbation during Learn trials [64]. The DO model predicts that the motor command would be:

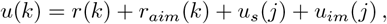

with *r*_*aim*_ = −*γ* · *d* and *γ* ∈ [0, 1] captures what fraction of the perturbation *d* the participant is informed about. Simulation results are shown in Figure 15 for three values of *γ*, and with *K* = 0.1 to accentuate the contribution of the aiming strategy. As *γ* is increased so that the motor command includes a larger explicit component, the learning rate increases, while the aftereffects decrease. We may compare these model predictions with the study in [64] that explored the relationship between reaction time, rate of decrease of errors, and aftereffects in a visuomotor reaching experiment with a perturbation of 60°. They found that a rapid decrease of errors was correlated with prolonged reaction times as well as reduced aftereffects. The participants with the longest reaction times exhibited the smallest aftereffects.

**Fig 15.**
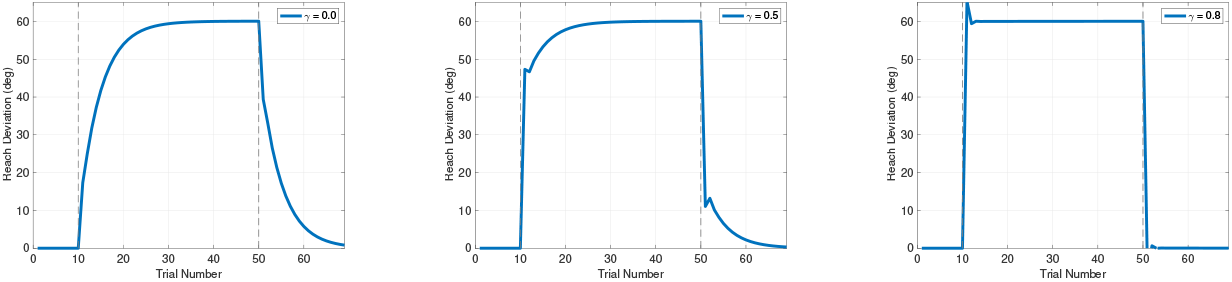
Feedforward strategy. We simulate a feedforward strategy *u*_*f*_ (*k*) = *r* −*γ* · *d* with *γ* ∈ {0, 0.5, 0.8} to emulate an aiming strategy in which participants are informed of and compensate for a fraction of the perturbation during Learn trials. As *γ* is increased so that the motor command includes a larger explicit component, the learning rate increases, while aftereffects decrease.

Longer reaction times are correlated with more emphasis on explicit strategies modeled by *u*_*exp*_(*k*). The DO model predicts that this component will reduce errors faster than a disturbance observer that operates on a (relatively) fixed learning rate determined by parameter *F*. Aftereffects arise in washout trials, and they correspond to unlearning in the disturbance observer of the previous 60° rotation. If the participant utilizes an explicit strategy, then the amount of learning to be performed by the disturbance observer is reduced, and this results in reduced aftereffects. The DO model does not capture any notion of time but we deduce from the discussions in [64] that explicit cognitive strategies take more time to constitute than the implicit component. See also the studies in [65] where further results (consistent with the study in [64]) on the role of explicit strategies are reported. A related and intriguing study is the one in [66] on decomposing explicit and implicit components of learning (with differing expressions of savings) as a function of preparation time.

### 5.7 Learning Transfer

Long-term adaptation has been described as a process that recalibrates motor skills, allowing the visuomotor system to adjust to changes in the body or environment without relearning from scratch [67]. In the study of [67] participants performed a visuomotor rotation task for one hour each day over five consecutive days. While participants demonstrated savings day to day, probing trials (in which participants indicated their intended aiming direction) revealed that participant’s aiming direction remained constrained. These findings complicate efforts to model long-term adaptation. The DO model offers a candidate mechanism for long-term adaptation - referred to in this context as learning transfer - that helps account for recent experimental results.

In extended visuomotor experiments, the DO model captures the emergence of long term adaptation through (13g). During Learn trials, this component of the motor command continues to grow, despite the fact that *x*_*f*_ (*k*) is limited due to the saturation imposed by *L*(*j*). The parameter *L*_*f*_ ≪ 1, so that (13g) represents a slow, long-term adaptation process. The combination of the short-term adaptation processes associated with *ŵ*(*j*) and *x*_*f*_ (*j*), combined with the long-term adaptation process for *r*_*f*_ (*j*) gives rise to a two timescale system. This should not be confused with a traditional two-rate model; rather it reflects a layered architecture in which long-term learning gradually consolidates alongside faster processes.

Figure 16 illustrates the DO model predictions for accelerated learning transfer in Experiments 1-4 with *α* = 1 and increasing values of *L*_*f*_, the rate of learning transfer. In Fig 16(a) ( Ignore-N Experiment) two primary effects are observed in the simulations: (i) an increase in the slope of the response during Ignore trials, and (ii) an increase in the steady state response in the no cursor trials. In Fig 16(b) (Learn-N Experiment) there is no discernible effect during the Learn trials, as these are governed by the disturbance observer. However, the effect of increased learning transfer is revealed during the no cursor trials (following a phase switch) where the steady state response increases with increased values of *L*_*f*_. Fig 16(c) (Ignore-W Experiment) shows that the slope of the response increases during the Ignore trials, but after a phase switch, no discernible effect is observed in the washout trials. In Fig 16(d) ( Learn-W Experiment) there is no discernible effect because the Learn and washout trials are both governed by the disturbance observer, leaving the learning transfer process unexpressed.

**Fig 16.**
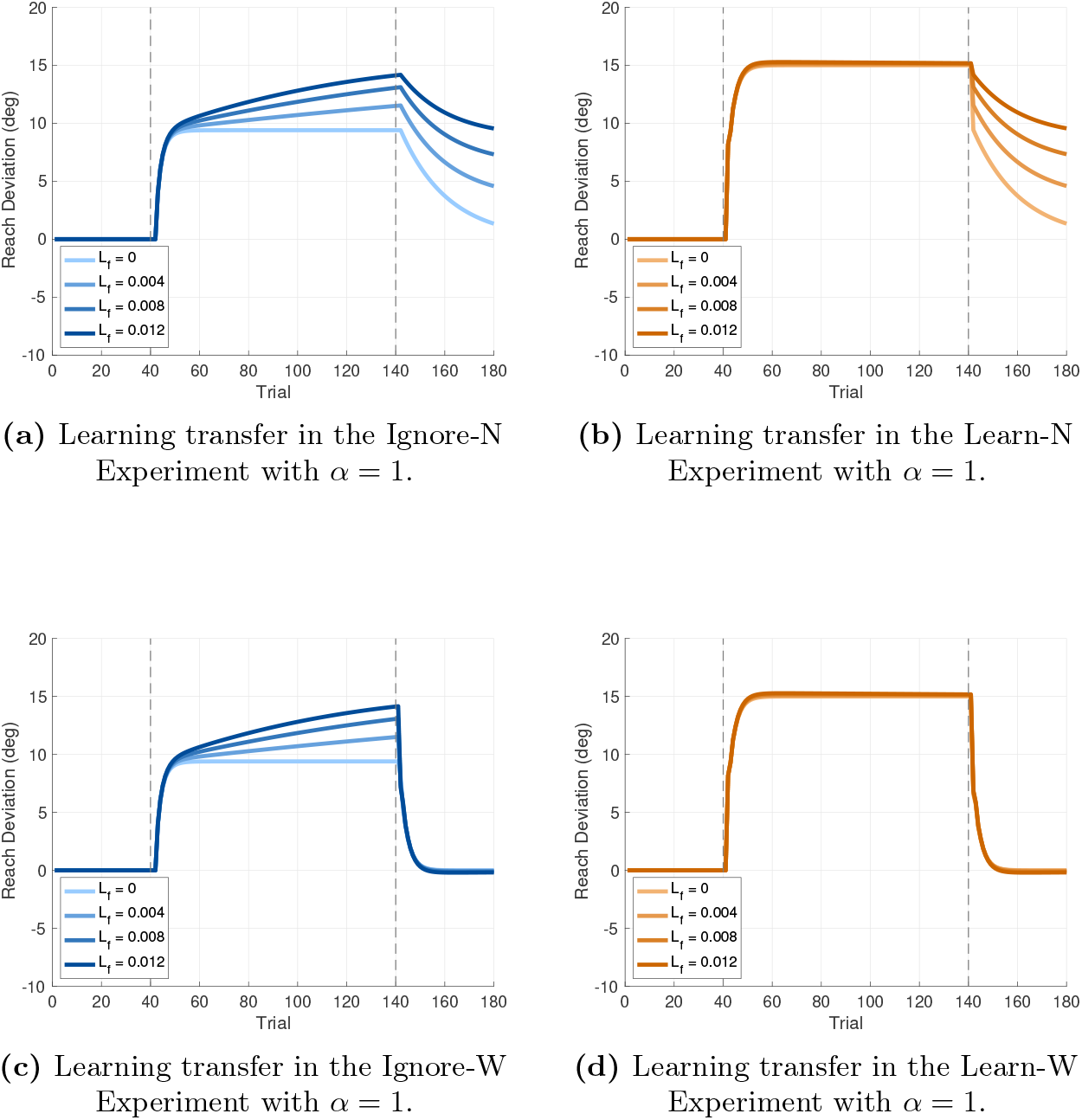
Learning transfer. Predicted behavior in Experiments 1-4 with *α* = 1 as a function of the learning transfer rate *L*_*f*_. All traces correspond to model predictions.

We next examine experimental data for evidence supporting the DO model’s predictions of learning transfer. Three datasets are potential candidates: the Ignore-N Experiment of the present study, the study reported in [18], and the study in [38]. Signs of learning transfer are present in all responses of Experiment 1, with the upward slope during Ignore trials particularly evident in Fig 9(c) with *α* = 0.6. Additionally, the steady state hand angle during no cursor trials is non-zero, both for Experiments 1 and 2, consistent with the DO model. An exception is found in the Learn-N Experiment with *α* = 0.8, where the hand angle returns to zero. Experiments 3-4 also exhibit evidence of learning transfer during Ignore trials, especially for *α* = 0.5, as seen in Fig 11(c). Indeed, smaller *α* may induce stronger learning transfer. The study reported in [18], illustrated in Fig 13, reveals additional behaviors consistent with learning transfer, which we examine next.

The DO model predicts that learning transfer will produce a non-zero slope during both cursor and no cursor trials. It also predicts the emergence of a bias in the final phase of zero error clamp trials - a bias that unlike spontaneous recovery (which is transient) behavior) reflects a stable shift driven by learning transfer. According to the DO model, this bias will not develop in the absence of learning transfer. Finally, the study in [38] reveals yet another aspect of learning transfer. Here the DO model predicts that the slope of the response during no cursor (Ignore) trials increases with perturbation size - an effect observed in the experimental data in Fig 14. Taken together, these three experiments demonstrate that learning transfer can emerge within a single experimental session; it does not require days of experiments. It is possible that paradigms such as those used in [18] and [38] accelerate the transfer process through rapid phase switching between two computational processes that exchange information.

Although experimental evidence suggests that learning transfer can develop over a single session, the use of a two timescale architecture (with a small value of *L*_*f*_) remains important for ensuring the stability of the system. Notably, the disturbance observer output *u*_*im*_(*j*) drives the update of the feedforward state *x*_*f*_ (*k*) in (13j), which in turn drives the adaptation of *r*_*f*_ (*k*) in (13g). However, *r*_*f*_ (*k*) modulates the update of *ŵ*(*j*), as computed in (8), thereby forming a closed feedback loop. If *r*_*f*_ (*k*) were updated on the same timescale as *ŵ*(*j*) and *x*_*f*_ (*k*), then this closed-loop system could potentially introduce instability and generate runaway behavior. The slower adaptation rate governed by *L*_*f*_ thus serves as a stabilizing mechanism within the system.

## 6 Discussion

Researchers have long debated that multiple processes contribute to visuomotor adaptation [68]. This paper proposes a model composed of two interacting components. The primary mechanism is a disturbance observer that builds an internal model of persistent exogenous disturbance and reference signals acting on the visuomotor system based on visual error. A secondary mechanism - a feedforward system - learns from the output of the disturbance observer and is responsible for generating motor commands that position the hand at a specific spatial location. Functionally, the feedforward system plays a key role in offloading the steady state demands on the disturbance observer. We hypothesize that the disturbance observer is biologically expensive to operate over prolonged periods, and that the feedforward system serves to reduce this cost by gradually assuming control of learned behavior. This dynamic contributes to a division of labor between short-term, error-driven correction and longer-term consolidation.

The architecture of the DO model is characterized by a cascade structure: the disturbance observer drives the feedforward system, and sensory signals do not act directly on the feedforward system. This separation is crucial for reproducing several known behaviors in visuomotor learning. For example, it aligns with findings from [69] which showed that error sensitivity scales with perturbation consistency during Learn trials but not in Ignore trials - suggesting different computational pathways underlie these contexts, as captured in our model. Another key feature of the DO model is the concept of *phase switches* - transitions in the dominant computational process that occur in response to changes in task structure or sensory input. Here, we focused on phase switches triggered by instruction changes (e.g., from moving the cursor to the target to moving the hand directly) and by changes in visual feedback (e.g. removal or reintroduction of the cursor). These phase switches allow for context-dependent engagement of different modules within the visuomotor system. A more comprehensive classification of phase switch types - potentially involving implicit contextual cues, motor plan changes, or uncertainty - would be a valuable direction for future research.

One of the model’s key predictions is that learning transfer is a meaningful and measurable phenomenon in visuomotor adaptation, with a distinctive behavioral signature. Specifically, it produces a gradual upward slope in hand angle during Ignore trials and a steady state bias in subsequent no cursor or zero error clamp trials. This dissociation underscores the importance of using trial structures that can selectively engage or reveal the contribution of the feedforward system - particularly when probing long-term or latent forms of adaptation.

### 6.1 DO Model Performance

Table 3 summarizes the parameter values used in the simulations. These values were not optimized in any formal sense; rather we selected nominal values that were sufficient to reproduce a wide range of qualitative behaviors across experiments. Notably, the parameters associated with the disturbance observer were remarkably stable across simulations. Slight variability was observed in the learning rate parameter *F*, which may reflect differences in experimental design - for example, the number of targets used, whether hand angles were averaged over targets, the use of continuous versus endpoint cursor feedback, or the duration of cursor visibility at the end of a reach. Parameter *ψ*_o_ was used as a proxy for direction-dependent biases in learning. Specifically, we used *ψ*_o_ = 0.8 for CW perturbations, *ψ*_o_ = 1.0 for CCW perturbations, and *ψ*_o_ = 0.9 when averaging across both. These minor adjustments were sufficient to account for incomplete adaptation effects and yielded consistent fits across studies. In contrast, greater variability was observed in the parameters associated with the feedforward system, particularly *A*_*f*_ and *b*_*f*_. The variability in *b*_*f*_ mirrors the variability in the steady state behaviors reported for error clamp experiments [22, 23, 25], suggesting it may be tied to experimental protocols. Whether a canonical value of *b*_*f*_ exists remains an open question and may be resolved through additional experimental studies. The variability in *A*_*f*_, however, points to a deeper issue: either the feedforward system is inherently more labile than the disturbance observer, or it reflects behavior that is not yet fully captured by the current DO model.

**Table 3.** Model parameters for simulations. Entries with a * indicate the parameter plays no role in the behavior. Note that parameter *F*_*n*_ is only discussed in Section 6.1.

| Behavior | Experiment | $F$ | $F_n$ | $\psi_o$ | $A_f$ | $A_{fn}$ | $b_f$ | $L_f$ |
| --- | --- | --- | --- | --- | --- | --- | --- | --- |
| Classical Behaviors | Savings with CP | 0.9 | * | 1 | * | * | * | * |
|  | Spontaneous Recovery | 0.9 | * | 1 | * | * | * | * |
|  | Evoked Recovery | 0.9 | * | 1 | * | 0.95 | 0.05 | * |
|  | Bansal, 2023 | 0.7 | * | 0.9 | 0.9 | 0.95 | 0.05 | * |
|  | Bond & Taylor, 2015 | 0.7 | * | 1 | * | 0.95 | 0.05 | * |
| Error Clamp | Morehead, 2017 | 0.7 | * | * | 0.9 | 0.95 | 0.05 | * |
|  | Kim, 2018 | 0.7 | * | * | 0.9 | 0.95 | 0.05 | * |
|  | Tsay, 2021 | 0.7 | * | * | 0.95 | 0.95 | 0.1 | 0.005 |
|  | Ignore-N (CW) | 0.7 | * | 1 | 0 | 0.95 | 0.05 | 0.005 |
|  | Ignore-N (CCW) | 0.7 | * | 0.8 | 0 | 0.95 | 0.05 | 0.005 |
|  | Learn-N (CW) | 0.7 | * | 1 | 0 | 0.95 | 0.05 | 0.005 |
|  | Learn-N (CCW) | 0.7 | * | 0.8 | 0 | 0.95 | 0.05 | 0.005 |
|  | Ignore-W | 0.7 | * | 0.9 | 0 | * | 0.01 | 0.002 |
|  | Learn-W | 0.7 | * | 0.9 | 0 | * | 0.01 | 0.002 |
| Savings | Avraham, 2021 | 0.9 | 0.92 | 0.9 | 0.9 | 0.95 | 0.05 | * |
|  | Fig 2A, Yin, 2020 | 0.85 | 0.95 | 0.9 | 0.6 | 0.92 | 0.02 | * |
|  | Fig 2B, Yin, 2020 | 0.85 | 0.9 | 0.9 | 0.6 | 0.92 | 0.05 | * |
| Phase Switch | Ruttle, 2021 | 0.7 | * | 1 | 0 | 0.8 | 0.01 | 0.005 |
|  | Amario, 2024 | 0.7 | * | 1 | 0 | 0.9 | 0.01 | 0.005 |

Table 3 also lists the experiments considered in this study, arranged roughly in order of increasing complexity from a modeling perspective. Classical behaviors such as savings, anterograde interference, and spontaneous recovery are governed by relatively few parameters and can often be reproduced using a standard two-rate model [44]. Evoked Recovery [45], the Bansal study [52], and the Bond and Taylor experiment [62] introduce additional complexity by probing aftereffects in no cursor or zero error clamp trials. These three experiments are predicted to involve a phase switch, whereby the parameters of the feedforward system play a central role in determining the magnitude of aftereffect or spontaneous recovery. The experiments listed near the bottom of Table 3 represent the highest level of complexity, as they require activation of all components of the DO model. In particular, the studies in [56] and [58] necessitated the introduction of a forgetting factor *F*_*n*_ in the disturbance observer state *w*_0_(*k*) during no cursor trials. Specifically, the update rule *w*_0_(*k* + 1) = *F*_*n*_*w*_0_(*k*)) was applied to account for apparent decay in the perturbation estimate. However, we chose not to make this a permanent feature of the model, as these experiments likely involve multiple phase switches, the precise number and timing of which remain uncertain. We return to this issue next.

An important consideration when evaluating the DO model performance is the possibility that more phase switches occur than currently identified; see [54] for a related discussion. For example, in the study in [52] spontaneous recovery was observed in the zero error clamp trials following a learning phase and short unlearning phase. This spontaneous recovery can be explained in one of two distinct ways. One interpretation attributes spontaneous recovery to a form of memory retention within the feedforward system. Setting *A*_*f*_ = 0.6 to 0.9 allows the feedforward system to retain part of the learned response across the unlearning phase. The parameter *b*_*f*_ modulates the degree of saturation, shaping the amplitude of recovered responses. Thus, spontaneous recovery during the zero error clamp phase can be predicted by the interplay of two parameters: *A*_*f*_, controlling retention, and *b*_*f*_, controlling saturation.

An alternative explanation assumes no memory in the feedforward system so *A*_*f*_ = 0, and instead posits that spontaneous recovery arises from a phase switch between distinct internal models. Under this view, the short unlearning phase engages a second feedforward system that learns the reverse perturbation. When the zero error clamp trials are introduced, a phase switch returns the computations to the feedforward system of the first internal model. The change in error direction may serve as a trigger for this switch. Given the present state of knowledge, we assume that only a single internal model is active in this experiment. However, further studies are needed to determine whether multiple feedforward systems operate concurrently or sequentially in such paradigms, and to clarify the mechanisms that drive phase switching under these conditions.

One experiment among the group we examined where the DO model does not make correct predictions is the study of [18]. We see in Figs 13(b)-(c) that during cursor trials with a counter-perturbation, the DO model does not match the response. The pattern of mismatch is the same pattern as in the Learn-N Experiment for CW v.s. CCW perturbations. To allow the model to match the experimental data, we may rescale the disturbance observer response using the parameter *ψ*_o_ for the counter-perturbation phase. This phenomenon raises the question of whether the observed behavior reflects a phase switch to a different internal model that is not presently captured by the DO model. Thus, there would be two separate internal models to handle CW or CCW perturbations.

### 6.2 Alternative Modeling Approaches

To motivate the modeling decisions that lead to the DO model, we discuss alternative modeling approaches falling into four categories: (i) approaches that violate the internal model principle; (ii) approaches we tried but did not work; (iii) approaches which we do not recommend at present, but may be relevant in future studies; (iv) alternative designs for the internal model and the feedforward system.

A crucial point to emphasize is that we only study visuomotor experiments in which learning takes place with constant perturbations arising in the visual error. As far as we are aware, there is no experimental evidence suggesting that the brain is capable to learn any more complex perturbation during short-term visuomotor adaptation than this. By *learn a perturbation*, we mean that mathematically, the perturbation can be completely eliminated from the visual error in steady-state. In fact, experiments have shown that the brain cannot even learn a sinusoidal visuomotor perturbation, since the steady-state error is never driven to zero [32]. The DO model is based on the premise that the human visuomotor system is crafted to reject predictable exogenous signals of low order. This is not so much a limitation of our model, but rather reflects what appears experimentally to be an inherent limit of the system.

The first major class of alternative modeling approaches one might consider would ignore the internal model principle of control theory, meaning the model would omit an internal model of the environment as part of the brain’s computations. It might utilize an internal model of the plant, an observer of the plant states, or several other constructs from control theory. On theoretical grounds, such a model will fail to capture the fact that the visual error can be driven to zero by the brain using solely the error itself as a measurement. For an error-driven process, the inclusion of an internal model of the environment determines a hard boundary on performance. For this reason, the disturbance observer is the centerpiece of the DO model. It provides the needed internal model of the perturbation that ensures the visuomotor system can perform with high precision using limited measurements.

A second modeling approach would place more emphasis on the plant. The focal point would be on the dynamics of the arm and a corresponding internal model of the arm in the brain. This approach is not recommended for two reasons. First, visuomotor adaptation is not a problem of mismatched parameters between a physical system and its internal representation. It regards the appearance of an unexpected perturbation in a visual error measurement, a signal with a fleeting lifespan. Re-calibrating an internal model of the arm in the presence of perturbations that come and go as tasks change is highly inefficient, given the complex computations needed to emulate the nonlinear kinematics and dynamics of the arm. A more biologically efficient approach would be to carefully guard internal representations of the body from corruption by exogenous factors until sufficient time has passed that such exogenous factors are deemed worthy to be internalized. As such, we do not see the modification of the brain’s internal model of the arm as the central problem in (short-term) visuomotor adaptation. A more intractable problem is that if we focused the model in modification of parameters of an internal model of the arm, then the internal model principle of control theory would be violated. In simple terms, an internal model of the arm dynamics is incapable to resolve the presence of a perturbation in a visual error signal.

A third modeling approach would place more emphasis on parameter adaptation laws. One may endow parameters in the model with parameter adaptation laws corresponding to the gradient law (in discrete-time, the Widrow-Hoff rule). The gradient law is a nonlinear update that is well known in control applications to generate wild transients. These extra transients create unreasonable responses which have no resemblance to the responses in visuomotor adaptation [37]. In practice, to tame the transients of the gradient law, one must use a high gain on the error feedback and low learning rates on the adaptation laws (as is done in robotics [24]). High gain on the error feedback wipes out the correct transient response by creating overshoot. A better method is to use a small learning rate on the gradient law, making the model parameters quasi-static over the time period of a typical short-term visuomotor experiment. We come full circle with the presented model where the parameters are constants. A further argument for not including parameter adaptation in the present model is that the responses in the visuomotor experiments we examined, with the exception of the error clamp, are manifestly linear responses; i.e. responses arising from linear state space models.

We have argued that the internal model principle of control theory must be accounted for in any plausible model of visuomotor adaptation. Further, there is little motivation to include detailed models of the plant or nonlinear parameter adaptation laws at this stage of model development (though, as more refined models appear, these extensions can be considered). It remains to describe our modeling choices for the feedforward system and the disturbance observer.

The feedforward model presented in the paper is not the first one we considered. Originally we attempted to utilize a standard Luenberger observer of the plant along with the disturbance observer modeling the environment. We were unable to recover all experimental results using this approach. Potentially a different observer design than the Luenberger observer may be needed, but we did not further pursue that avenue. The current feedforward model was inspired by prior work on modeling the floccular complex (FC) [15, 16]. The floccular complex model uses an internal model based on [70], and it includes a feedforward process to capture learning in the medial vestibular nucleus based on the Purkinje cell output of the FC. This control architecture was the key inspiration for the current model of the feedforward system. It also informs why the control architecture of the DO model has a dominantly cascade structure whereby the disturbance observer trains the feedforward system.

Finally, we motivate our design choice to use the disturbance observer from [31]. First, consider the classical internal model of [36]:

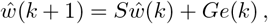

where *e*(*k*) is any scalar error signal being regulated to zero, *ŵ*(*k*) ∈ ℝ^*q*^ is the state of the internal model, and *S* ∈ ℝ^*q×q*^ is a matrix whose eigenvalues include all modes present in the disturbance being rejected (for a precise mathematical statement of the requirements on *S*, see [36]). This design is reasonable for classical unity feedback loops in which the error is a permanently available measurement (a biological example is blood sugar regulation). In visuomotor adaptation, the error *e*(*j*) disappears in No Cursor trials, or it simply becomes irrelevant when the visuomotor task ends.

Unfortunately, there is no clear mechanism to “turn off” the classical internal model - indeed, it was never conceived to operate this way. Removal of the error signal from the internal model does not suffice, because the eigenvalues of *S* are marginally stable for persistent signals, so *ŵ*(*k* + 1) = *Sŵ*(*k*) produces a steady-state drive. Such a model cannot be housed in the brain, where mechanisms to turn on and off internal models are needed for safe operation.

Second, consider the internal model of [70]:

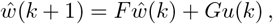

where *F* ∈ ℝ^*q×q*^ is a Schur stable matrix, *ŵ*(*k*) ∈ ℝ^*q*^ is the state of the internal model, and *u*(*k*) is the motor command. This internal model improves on the classical one. It includes a recurrent architecture in which the output of the internal model *u*_*im*_(*k*) is included as an input to the internal model using

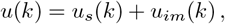

where *u*_*s*_(*k*) is an error feedback for stability. This internal model has a clear mechanism to activate or de-activate itself: the recurrent loop can be disabled by removing the input *u*(*k*), such that the remaining dynamics *ŵ*(*k* + 1) = *Fŵ*(*k*) are stable. As such, this internal model is a candidate for study of Learn and No Cursor trials. Unfortunately, it does not function properly for Ignore trials. During Ignore trials, the subject performs a different task, move the hand to the target, meaning the wrong motor command will be supplied to the internal model.

The disturbance observer we have selected has the outstanding property that it can learn the perturbation even when the subject is instructed to ignore the cursor; in fact, it is agnostic to the visuomotor task. In mathematical terms, the update of *ŵ*(*k*) does not depend on the value of *u*(*k*), as verified in the computation (8).

### 6.3 Model of the Saccadic System

The DO model is targeted to modeling behaviors during visuomotor experiments with constant perturbations in the visual error. To apply the model to other scenarios such as the saccadic system or force field experiments, it is necessary to adjust the model to capture the physics of the new experiment. For didactic purposes, we demonstrate how such an adjustment can be made to model the saccadic system in *intersaccadic step* (ISS) experiments [43].

The state *x*(*k*) represents the saccade amplitude on the *k*-th saccade; the desired saccade amplitude on the *k*-th trial is given by *r*(*k*) + *d*(*k*); *r*(*k*) denotes the nominal saccade size based on the location of a visual target relative to the initial eye position; and *d*(*k*) represents a perturbation introduced by the experimenter while the *k*-th saccade is in progress. The visual error experienced by the participant at the end of the *k*-th saccade is therefore

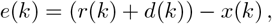

representing the mismatch between the required and actual saccade amplitude.

The saccadic system operates through *adaptation fields* - spatially localized, and often overlapping regions on the retina, each governed by its adaptation process. To isolate and model a single adaptation process, the range of saccade magnitudes *r*(*k*) should be constrained to fall within a single adaptation field. For simplicity, we assume a constant saccade amplitude within this range such that *r*(*k*) ≡ *r*. The motor command for a given adaptation field includes a feedforward component that generates the appropriate movement for the nominal saccade size *r*. Thus, we set the feedforward term to be *u*_*f*_ (*k*) = *r*.

Collecting these adjustments, we arrive at a simplified model of the saccadic system for an adaptation field with a nominal saccade size *r*:

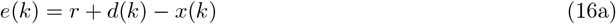

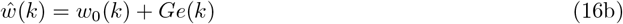

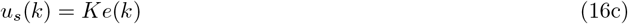

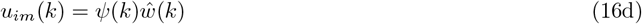

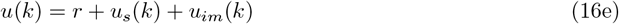

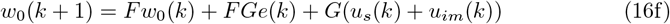

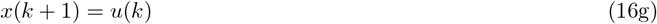

The term *u*_*im*_(*k*) captures short-term adaptive changes via a disturbance observer. The feedforward component *r*_*f*_ (*k*) = *r* remains fixed for the adaptation field. To simulate this model, we use the following nominal parameters values: *K* = 0.22, *F* = 0.9, *G* = 1 − *F*, and *ψ*_o_ = 0.95. Simulations of (16) reproduce a range of hallmark behaviors observed in visuomotor adaptation: savings, reduced savings, anterograde interference, spontaneous recovery, rapid unlearning, and rapid downscaling [37].

### 6.4 Comparison to Existing Models

#### 6.4.1 Two-rate Model

Arguably the most influential model of sensorimotor adaptation is the two-rate model of [44]. This simple two state framework has proven remarkably effective in capturing a wide range of adaptive behaviors. Here we delve into the reason why this model is so resilient.

Let us assume that the perturbation is constant, *d*(*k*) ≡ *d*, and *ψ*(*k*) ≡ *ψ*_o_. We identify the visual error *e*(*k*) as the fast state, and the disturbance observer estimate *ŵ*(*k*) as the slow state. Using this mapping, we can compute the updates for *ŵ*(*k* + 1) and *e*(*k* + 1) using the preliminary visuomotor model (10) to obtain the reduced model (11) (thus, we omit the feedforward system). The first equation is an error model that captures the evolution of the error. This equation explicitly includes the perturbation *d*, to reflect the fact that the perturbation acts externally on the system, shaping the error dynamics. These dynamics are clearly not internal to the brain; rather they unfold in the physical world but are observable as a measurement in the brain. The second equation captures the evolution of a brain state: the slow state *ŵ*(*k*) which gradually estimates the perturbation by way of a simple filter operation. In the DO model this internal estimation is achieved using both the observed error and an efference copy of the motor command.

While the system described in (11) captures core features of sensorimotor learning, it is not mathematically identical to the two-rate model in [44]. This highlights the point that the representation of learning processes depends on the choice of state variables, and that choice is not unique. The canonical two-rate model is typically written in the form

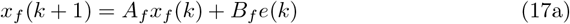

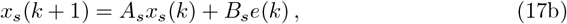

where *x*_*f*_ (*k*) and *x*_*s*_(*k*) denote fast and slow internal states, respectively, and the motor output *x*(*k*) is the sum of these components. Both processes are driven by the same error signal and act in parallel to produce behavior.

In this formulation, the slow and fast states are typically treated as abstract brain states, without assigning them specific physiological or anatomical meaning. An important feature of (17b) is that the parameter *A*_*s*_ is always set to a value close to 1, reflecting slow retention dynamics. This choice is significant as it aligns with the internal model principle of control theory [71, 72]. In this context the principle asserts that that for the brain to learn and compensate for a constant perturbation *d* using only error feedback, it must contain an internal model capable of generating constant signals. In control-theoretic terms, such a structure is known as an exosystem, and its canonical form is given by

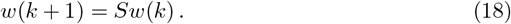

For this exosystem to generate constant signals, it is required that *S* = 1. In the two-rate model, this means *A*_*s*_ must be close to 1 in order to achieve an internal model of this exosystem. Equation (17b) is precisely the classical internal model introduced by E.J. Davison [36, Eq. 10], here for discrete-time systems. From a control theory perspective, the enduring success of the two-rate model lies in its ability to capture the fundamental - and mathematically necessary - requirement for an error-driven process to achieve perfect regulation in response to constant disturbances.

#### 6.4.2 Memory of Errors Model

The memory of errors (MoE) model, introduced in [73], is based on the principle that the brain can dynamically adjust the sensitivity to perceived errors. In this framework, persistent errors are assigned greater weight than variable or inconsistent errors, allowing the MoE model to explain phenomena such as saturation on single-trial learning [27]. The DO model does not include trial to trial modulation of error sensitivity. Instead, we assume a fixed sensitivity parameter *K* in the error feedback *u*_*s*_(*j*) = *Ke*(*j*). To remain within the linear range of this error sensitivity function, we deliberately chose small perturbation sizes in Experiments 1-4. Nonetheless, the structure of (13d) can be naturally extended to include saturation effects by allowing the error sensitivity parameter *K* to vary as a function of recent error history or magnitude. Such an extension could allow the DO model to account for trial-level nonlinearities and adapt more closely to the MoE framework, while preserving the architectural separation between disturbance estimation and feedforward adaptation.

The DO model (13) also incorporates two error sensitivity functions defined in (15a)-(15b), which bear conceptual similarity to the error sensitivity mechanisms proposed in the MoE model. A key distinction, however, is that the MoE model attributes error sensitivity to a population coding. In contrast, the DO model employs static error sensitivity functions that depend on the instantaneous error; they do not incorporate any temporal integration or memory of past errors. At this stage it remains unclear whether one approach offers a fundamentally more accurate description of neural error processing. While both frameworks can replicate key features of motor adaptation behavior, their underlying mechanisms differ significantly. It is likely that future neurophysiological investigations - particularly those examining how error sensitivity is represented and modulated across different brain regions - will be required to adjudicate between these two accounts or to motivate a hybrid framework.

Finally, the MoE model also includes an explicit model of the environment, taking the form

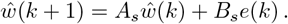

As in previous models, the inclusion of an internal model within the MoE framework is necessary to satisfy the internal model principle. Specifically, accurate compensation for constant perturbations depends on incorporating an internal representation of the perturbation dynamics. This requirement is formalized by setting the adaptation parameter *A*_*s*_ close to or equal to 1. Only under this condition can the system achieve correct asymptotic behavior in response to constant disturbances.

#### 6.4.3 COIN Model

The COIN model, introduced in [45], offers a distinct framework for understanding sensorimotor adaptation. In this model, each context has associated with it a scalar linear state equation that models a latent environmental quantity - such as the imposed perturbation in a visuomotor task. This formulation allows the model to maintain and update an arbitrarily large number of context-specific internal models. For each context, a corresponding observation provides a noisy replica of the context state, while a Kalman filter is employed to produce a noise-free estimate of the context state. The motor command is then computed as a weighted sum of these state estimates, where the weights correspond to the inferred probabilities (or responsibilities) of each context. Importantly, the model updates all contexts at each time step, scaling the updates by their respective inferred responsibilities.

There are several points of comparison between the DO model (13) and the COIN model. A point of commonality is that both models posit that multiple neural processes are updated in parallel and both allow for *silent* updates - i.e. changes in internal states that may not be immediately reflected in behavior. However, the models diverge in important ways. The COIN model is grounded in probabilistic reasoning for maintaining contexts, each associated with its own Kalman filter estimate and context probability. In contrast, the DO model operates deterministically and assumes a binary instructional context - whether the participant is instructed to move the hand or the cursor to the target - eliminating the need for probabilistic context inference.

A more profound difference between the DO model (13) and the COIN model regards the measurement structure. The COIN model assumes that the brain receives a noisy observation of each latent context state, enabling Kalman filters to denoise the signal. For the visuomotor experiment, this implies the brain receives a noisy measurement of *d*(*k*). The COIN model does not explain how the the brain obtains access to this signal. The DO model, on the other hand, utilizes a single measurement *e*(*k*) to reconstruct the perturbation, showing how the brain can estimate an inaccessible exogenous signal.

Another conceptual innovation in the DO model is the notion of a phase switch, which is a switch in the computational process governing motor output. This is distinct from a context switch, which refers to a change of external task conditions or instructions. For example, while a transition from Learn to washout trials represents a context switch, the underlying computations may remain unchanged - no phase switch occurs. Instead the transition from Learn to No Cursor is both a context switch and a phase switch according to the model. Identifying and classifying the computational phase switches that occur during visuomotor adaptation is an important direction for future research.

#### 6.4.4 CPC Model

A recently proposed framework is the cerebellar population coding (CPC) model [74] offering another biologically grounded approach to modeling sensorimotor adaptation. The CPC model is inspired by the Marr-Albus model of the cerebellum, but it also includes a model of plasticity in the deep cerebellar nuclei (DCN). The cerebellar cortex is hypothesized to act as an internal model to predict the sensory consequences of motor commands, positioning the cerebellum as either a forward or inverse model. The model adheres to cerebellar architecture in which Purkinje cell outputs project to the DCN.

A key architectural similarity between the DO model and the CPC model lies in their hierarchical (or cascade) structure. In the DO model, the feedforward system is driven by the disturbance observer, which is hypothesized to be implemented in the cerebellum. This mirrors the known anatomical pathway in which Purkinje cells project to the deep cerebellar nuclei (DCN), aligning with the CPC model’s mapping of the cerebellar cortex and DCN roles. However, the DO model is a behavioral-level framework and does not place a restriction that the feedforward system resides entirely within the DCN. It may instead involve additional downstream structures, such as cortical motor areas. In this sense, the DO model maintains a more abstract relationship to cerebellar anatomy than the CPC model, which aims to more closely reflect the underlying neural circuitry. This distinction echoes the contrast between phenomenological and biophysical models of synaptic plasticity — where the former aim to capture functional behavior and the latter emphasize mechanistic fidelity.

A key distinction between the DO model and the CPC model lies in the interpretation of the visuomotor system’s core computational function - particularly in regard to the cerebellum. We propose that the primary function of the cerebellum is to build internal models of persistent exogenous reference and disturbance signals in the environment, in line with the internal model principle of control theory [71, 72]. In contrast, the CPC model assumes the cerebellum builds forward (or inverse) models (putatively of the plant), thus estimating the current state of the effector (e.g. hand angle) given the motor command. In terms of expressive capability, the CPC model focuses exclusively on behaviors observed during error clamp conditions. The DO model is designed to capture a broader range of behaviors in visuomotor adaptation, including classical savings and spontaneous recovery, as well as more complex behaviors involving phase switches and long-term learning transfer. In particular, the CPC model does not a priori account for the results of Experiments 1-4, which go beyond a strict error clamp paradigm.

#### 6.4.5 Summary of Comparison

We summarize the comparison with other models, in terms of these metrics: (i) adherence to the internal model principle of control theory; (ii) biologically plausibility of the measurement structure; (iii) suitability of the control architecture; (iv) ability to capture learning transfer and phase switches; and (v) low-dimensional models. A metric not included in this study is numerical performance, and this is because the primary contribution of the DO model is improved understanding, not improved performance. Additionally, performance comparisons may raise questions of soundness of the comparison when dealing with models with vastly different measurement structures and model orders, etc. Nevertheless, some remarks on expected performance differences can be helpful.

For experiments that involve standard learning which require only a disturbance observer (no feedforward system), meaning the cursor is always visible and there is no error clamp, the two-rate model will have equivalent or better performance than our model. This is because the two-rate model is an abstract model with no a priori limits on the values of its four parameters. The DO model, for the equivalent behavior, has three free parameters *K, F* and *ψ*_o_. In practice, *K* has little variability, so only two parameters should be assigned. Parameter *F* can only be used for the learning rate in Learn trials, and *ψ*_o_ is the sole parameter that sets the proportion of learning at asymptote. Thus, the selection of parameters in the DO model is severely restricted under proper usage. The DO model classifies the slow state *ŵ*(*k*) as a brain state and the fast state *e*(*k*) as measurable by the brain but evolving in the world, while the two-rate model blurs this boundary by considering only abstract states. The resilience of the two-rate model arises from its adherence (with a suitable parameter assignment) to the internal model principle; however, it was not conceived to capture learning transfer or phase switches.

The MoM model, as a higher-order extension of the two-rate model, adheres to the internal model principle (via a so-called memory of the state of the environment), well explains saturation in single-trial learning and certain types of savings, but does not capture learning transfer. We submit the latter behavior emerges strictly from a cascade architecture in which the error only enters the computations via the disturbance observer, not the feedforward system. A performance comparison with the memory of errors model may be deceiving. Each past value of the error that is stored by the MoM model corresponds to an extension of the state vector. As such, the MoM model can be significantly higher order than the DO model, which works with only three controller states. Using performance alone, modelers may be erroneously led to choose a higher order model over a low order model, which bypasses considerations of biological cost.

The COIN model does not incorporate the internal model principle; consequently this model is forced to impose a biologically implausible assumption that a noisy measurement of the perturbation is already available in the brain. Instead, the DO model clarifies how a disturbance observer can reconstruct the perturbation using only sensory error and an efference copy of the motor command. The COIN model is targeted to explaining how motor plans are selected, so it offers a complementary modeling perspective to the DO model, which characterizes error driven processes to eliminate a perturbation in a visual measurement. A performance comparison between two models with different measurement structures should be avoided, as a clear interpretation of results may be elusive.

Finally, the CPC model is based on a Marr-Albus framework to implement forward or inverse models (typically driven by prediction errors). Since the CPC model focuses only on error clamp behavior with instructions to ignore the cursor, it does not incorporate phase switches due to varying instructions or due to removal or reintroduction of the cursor. Similarly, learning transfer, a process involving learning a perturbation during either Learn or Ignore trials, is not captured in the CPC model.

### 6.5 Biological Plausibility

We have presented a discrete-time model of visuomotor adaptation in reach tasks. Like all biological discrete-time models, the formulation is necessarily abstract. Nevertheless, it is reasonable to ask whether the DO model components are biologically plausible.

The proposed disturbance observer is inspired by the adaptive internal model used to describe computations in the floccular complex [15, 16]. While an abstract discrete-time model, it may serve as a computational blueprint for cerebellar contributions to visuomotor adaptation [75]. For instance, the stable filter in (13i) may represent filtering in the cerebellar granular layer. The signal *u*(*k*), which drives this filter, aggregates multiple classes of mossy fiber inputs, including the component *u*_*im*_(*j*) that reaches the cerebellar component via the nucleo-cortical pathway [76–79]. The visual error signal *e*(*j*), critical for adaptation, could arise from activity in regions such as the superior colliculus as well as the frontal eye fields of the frontal cortex. Although the DO model does not explicitly represent the climbing fiber input as well as synaptic plasticity at parallel fiber - Purkinje cell synapses, these elements are essential in cerebellar learning. Prior work has modeled adaptation at these synapses as a slower process [37]. Further research is needed to bridge differences between the internal model used for the floccular complex and one that supports visuomotor adaptation. The nonlinear error sensitivity in (15a) may reflect modulatory input from the nucleoolivary pathway, which is known to to suppress cerebellar activity [10, 80, 81]. The parameter *ψ*(*j*) may be interpreted as a form of error gating or safety control, limiting unstable behavior in the internal model. Biologically, this would require graded inhibition in the inferior olive [82–84].

At present, the anatomical substrate for the feedforward system remains less certain. Due to its simplicity in the DO model, we can only speculate at this point. One might assign this role to the dentate nucleus, analogous to the manner in which learning transfer occurs from the Purkinje cells of the floccular complex to the medial vestibular nucleus [9, 11, 14, 85]. Alternatively, downstream targets of the dentate nucleus—including primary motor cortex (M1), prefrontal cortex, and posterior parietal cortex—may also contribute to feedforward computations [86, 87].

## 7 Conclusion

In this project we set out to explain various phenomena in visuomotor adaptation using a disturbance observer based model, a relatively recent innovation in control theory. The resulting DO model has three main components: a fast reacting error feedback, a somewhat slower disturbance observer, and a feedforward system that learns from the disturbance observer. The motor command switches between the output of the disturbance observer and the feedforward system, depending on the instructions to the participant to move either the hand or the cursor to the target. This discrete time model is necessarily a simplification, but its development has been guided by biological principles and anatomical structure. Initial model parameters for the error feedback and disturbance observer were based on previous literature; the error feedback is derived from single-trial learning and the disturbance observer model has strong ties to the well-established two-rate model. A key feature of the DO model is its cascade architecture, in which the feedforward system does not directly sense the visual error, but rather it learns from the disturbance observer.

Using the DO model we can explain several forms of spontaneous recovery, several types of savings, error clamp behavior, behaviors arising from switching between instructions, and explicit and implicit computations. We also expose several new behaviors. We introduce a novel experimental paradigm called the graded error clamp to explore the emergence of error clamp behavior over a sequence of experiments, by varying a single parameter that scales the hand angle. We introduce the notion of a phase switch, and we identify several forms of phase switches, based on current datasets. We introduce learning transfer in visuomotor adaptation, and we fully characterize its signature behaviors. The DO model does not fully capture experiments involving both counterclockwise and clockwise rotations, indicating that an extension with separate disturbance observers for each direction may be required. Exploring this possibility is left for future work. The DO model compares well in terms of its expressive capabilities relative to existing models; nevertheless, we see it as a complementary tool for further study. It utilizes a control architecture that mimics the computations in the floccular complex and medial vestibular nucleus for the slow eye movement systems. Analogies between models of the slow eye movement systems and visuomotor adaptation may provide a forward step toward unveiling the computations of the cerebellum, though further research is needed to determine if this analogy is fruitful.

Most importantly, a disturbance observer framework offers a new perspective from control theory that should be considered in modeling visuomotor adaptation.

## Acknowledgments

We acknowledge the generous support of the Government of Canada’s New Frontiers in Research Fund (NFRF), (NFRFE-2021-00458).

## Author Contributions

**Conceptualization:** Mireille E. Broucke, Bernard M. ‘t Hart, Denise Y.P. Henriques, Jean-Jacques Orban de Xivry

**Data collection and analysis:** Gaurav Sharma

**Funding acquisition:** Mireille E. Broucke, Denise Y.P. Henriques

**Writing - original draft:** Mireille E. Broucke, Gaurav Sharma

**Writing - review & editing**: Mireille E. Broucke, Bernard M. ‘t Hart, Denise Y.P. Henriques, Jean-Jacques Orban de Xivry

